# DNA methylation around transcription start sites not globally associated with transcription in the grain of natural and synthetic hexaploid wheat

**DOI:** 10.1101/2025.10.08.680649

**Authors:** Meriem Banouh, Mamadou Dia Sow, Caroline Pont, Michael A. Seidel, Jerome Salse, Peter Civan

## Abstract

Epigenetic mechanisms including DNA methylation are assumed to play crucial roles in the maintenance of genome integrity, regulation of gene expression and development, and their increasing exploitation in breeding applications is anticipated. However, the relationship between DNA methylation and gene expression remains ambiguous and difficult to generalize. Here we explored the hypothesized causality between the level of transcription and cytosine methylation at the 5’ end of genes (around transcription start sites and start codons) in relation to whole-genome duplication in natural and synthetic allohexaploid wheat (*Triticum/Aegilops* complex). Using transcriptomes and a sequence capture protocol coupled with bisulfite sequencing, we observed sometimes significant, but overall very weak associations between gene expression and 5’ end methylation on a genome-wide scale. In synthetic wheat allohexaploids, global methylation differences between subgenomes are not triggered by the polyploidization, as the subgenome patterns are rather faithfully inherited from parents. A small number of genes differentially methylated between the parents and synthetics was consistently recovered in reciprocal synthetics and subsequent generations. Differences in transcription between homeologs are not clearly associated with 5’ end methylation in either natural or synthetic wheat. Overall, allopolyploidization triggers only minor methylation changes around transcription start sites and start codons of nascent wheat allopolyploids, and these are not statistically associated with differential expression. Although there is a measurable methylation difference between expressed and non-expressed genes, our results do not support the hypothesis that 5’ end DNA methylation is engaged in the regulation of gene expression in natural and synthetic wheat.

**SIGNIFICANCE STATEMENT:** While a’genome shock’ hypothesis predicts extensive transcriptomic and epigenetic reorganization after polyploidization, DNA methylation patterns around transcription start sites are generally undisturbed in nascent wheat allohexaploids. Although this stability might indicate importance for gene regulation, a clear relationship between DNA methylation and transcription was not observed either on a genome-wide scale, or among triads of homeologous genes.

## INTRODUCTION

DNA methylation commonly refers to the addition of a methyl group to the C5 position of cytosines in the DNA molecule. This covalent modification widespread in the plant genome and across species is crucial for maintaining genome integrity by silencing transposable elements (TEs) (Galindo-González et al., 2018), and is thought to be the main component of the epigenetic regulation of gene expression (Moore et al., 2013; Zhang and Zhu 2024). In plants, methylated cytosines (mCs) occur in any sequence context (i.e. on cytosines followed by any nucleotide), and have been shown to be involved in various biological processes, including germline and embryo development, parental imprinting, fruit ripening (Zhang and Zhu 2024; Gu et al., 2024) and stress responses (Dowen et al., 2012; Zheng et al., 2022).

The regulatory role of DNA methylation is exerted on at least two different levels. Firstly, DNA methylation affects chromatin states and plays an instrumental role in heterochromatin formation (Zhang and Zhu 2024), with general impacts on transcriptional levels across larger genomic regions. Together with other epigenetic modifications, namely the histone variants H3K9me2, H3K27me1 and H2A.W, mCs determine a repressive chromatin state to ensure stable silencing (Sequeira-Mendes et al., 2014; Jamge et al., 2023). Secondly, DNA methylation affects the binding of transcription factors (TF) and other proteins to DNA. A group of methyl-binding domain proteins binds to methylated CG-dense sequences and subsequently interacts with an array of additional proteins to regulate the silencing or activation of associated genes through modulation of the state of protein aggregation (Zhang and Zhu 2024). In Arabidopsis, most TFs are sensitive to the methylation state of TF recognition motifs *in vitro*, with 72% and 4.3% of tested TFs preferentially binding to unmethylated and methylated DNA motifs, respectively (O’Malley et al., 2016). These findings lead to an expectation that heavy cytosine methylation is associated with transcriptional silencing, and that promoters of actively transcribed genes should be unmethylated in most cases. Cytosine methylation patterns of TEs (Transposable Elements), which should remain transcriptionally silent in most if not all tissues to maintain genome integrity, are in agreement with this expectation. In all examined plant genomes, CpG and CHG methylation (H representing any of A, C or T), and usually also CHH methylation, is distinctly elevated in TEs compared to the surrounding regions (Niederhuth et al., 2016). However, the relationship between DNA methylation and transcription is more difficult to untangle in the case of non-TE transcripts. Constitutively unmethylated genes can have a very wide range of expression levels. On the other hand, genes characterized by TE-like methylation (i.e. high methylation levels in all sequence contexts) are often times silent, but some of them can be highly expressed in specific tissues (Zeng et al., 2024). In most species, highly expressed genes are characterized with high CpG methylation and low CHG/CHH methylation in their bodies, forming a pattern called gene-body methylation (gbM). Nonetheless, gbM genes have the widest range of expression levels in most species (Niederhuth et al., 2016), leading to doubts about the role of cytosine methylation in gene expression. Since the loss of gbM in a *met1* mutant of Arabidopsis (defective in DNA methylation maintenance) does not affect gbM gene expression patterns (Bewick et al., 2016), it has been suggested that gbM has a minimal or no effect on gene expression, but possibly has a’homeostatic’ role in preventing aberrant transcripts (Muyle and Gaut, 2019). Nonetheless, one general feature of DNA methylation in relation to transcription is recognized. Averaged across all genes, including gbM genes, CpG methylation shows a marked dip around transcription start sites (TSS), and to a lesser extent also around transcription termination sites (Niederhuth et al., 2016). For many species, including grasses *Zea mays* and *Brachypodium distachyon*, CpG methylation at TSS is associated with almost complete silencing (Niederhuth et al., 2016). It therefore appears that the vicinity of TSS needs to stay free of any CpG methylation in order to enable the initiation of transcription.

Bread wheat (*Triticum aestivum* L.) is a monocotyledonous crop with a large genome (14.6 Gb pseudomolecule-level assembly of Chinese Spring v2.1; Zhu et al., 2021). The large genome size can be attributed to the combined effect of high TE content (85% of the assembly) and recent polyploidizations. Bread wheat is an allohexaploid (2n = 6x = 42 chromosomes; genomic constitution AABBDD) that emerged approximately ∼10,000 years ago from hybridizations and subsequent polyploidization involving tetraploid *T. turgidum* (2n = 4x = 28; AABB) and diploid *Aegilops tauschii* (2n = 2x = 14; DD) (McFadden and Sears, 1946; Marcussen et al., 2014). Due to its economic significance and the status of a worldwide staple crop, substantial efforts have been directed to characterize the bread wheat genome (Alaux et al., 2018). However, much less attention has been paid to the epigenome and the methylome (the complete set of DNA methylation modifications) in particular. A few large-scale studies (Gardiner et al., 2015; Ramirez-Gonzales et al., 2018; Li et al., 2019a) have reported that a genome-wide overview of DNA methylation in wheat matches the general patterns known from other species. However, more in-depth analyses are in demand, as the epigenetic regulation of gene expression is of special interest for epibreeding applications (Kawakatsu and Ecker 2019; Varotto et al., 2022; Singh et al., 2021), but also in relation to fundamental questions on the evolution of polyploid species. While most genes of bread wheat are in three homeologous copies (i.e. syntenic paralogs located on the A, B and D subgenomes of the hexaploid), a significant fraction of them is silenced (Mutti et al., 2017; Ramirez-Gonzales et al., 2018; Takahagi et al., 2018; Banouh et al., 2023), leading to frequent situations where the total expression of a’triad’ (the set of three homeologous genes) is supplied by one or two copies. This phenomenon of homoeolog expression imbalance is potentially significant for both evolutionary adaptation (homeolog neo-and sub-functionalization) and crop performance related to the’fixed’ heterosis of allopolyploids (Chen 2010, Qin et al., 2021). It may also be related to the transcriptional reprogramming commonly observed in nascent polyploids (Qiu et al., 2020; Tayalé and Parisod 2013) and hypothesized to have an epigenetic basis (Matzke et al., 2015). DNA methylation changes have been observed to accompany polyploidization in multiple species (Li et al., 2019b; Rao et al., 2023; Jiang et al., 2021; Xiang et al., 2023), including *Triticeae* (Yaakov and Kashkush 2011; Kraitshtein et al., 2010; Yuan et al., 2020; Miao et al., 2024), but their genome-wide assessment in relation to gene expression in allohexaploid wheat is missing.

With these considerations, we developed a hybridization-based sequence capture design targeting the 5’ end region (centered on TSS or start codon - SC) of most high-confidence (HC) genes of the wheat genome for the purpose of methylome analysis *via* bisulfite sequencing (BSseq). We specifically hypothesize that cytosine methylation around TSSs is employed by plants as a transcriptional on-off switch, and we tested this hypothesis by comparing the BS-seq and transcriptomic data from developing grain. We particularly focus on DNA methylation patterns in the polyploid context, exploring the relationship between DNA methylation and homeolog expression imbalance, monitoring DNA methylation changes in nascent allohexaploid wheat triggered by the whole-genome duplication, and examining possible links between differential gene expression and differential DNA methylation. We discuss our findings in the context of the underwhelming genomic response to polyploidization in wheat, and the persistently obscure relationship between DNA methylation and gene expression.

## RESULTS

### Capture-BSseq efficiency

Assuming that DNA methylation of promoters and TSSs is important for the transcriptional level of genes, we aimed to capture the 5’ ends of all HC genes annotated in the assembly of the Chinese Spring wheat genome. These intended targets are centered around TSS in most cases (150 bp upstream and downstream), but our capture probes targeted SC (150 bp upstream and downstream) in HC genes that lack 5’UTR annotation. We analyzed the methylation in the developing grain of natural hexaploid wheat cultivar (*cv.*) Recital, tetraploid durum wheat *cv.*Langdon, two diploid *Ae. tauschii* genotypes and five synthetic allohexaploids (Table S1), targeting 121,909 genes (Table S2). The synthetics include polyploid progeny from reciprocal crosses and two successive generations (109xL-C2, 109xL-C4 and Lx109-C2), and synthetics derived from alternative parental genotypes (Jx87-S5 and Jx109-S5; BS-seq data from the tetraploid parent missing). With two biological replicates per sample, we sequenced 18 libraries containing between 25 and 33 million read pairs (2×150 bp; Table S3). The mapping efficiency (i.e. the percentage of uniquely mapped reads in respect to the raw reads count) ranged from 36% to 54%. Out of the uniquely mapped reads, 43% to 51% overlap with the intended targets comprising only 0.26% of the Chinese Spring assembly, which demonstrates strong target enrichment. Overall, 18%-26% of raw BS-seq reads were used in the methylomic analysis (reads uniquely mapped to the target space). The bisulfite conversion rates estimated from the chloroplast genome-mapping reads exceeded 96% (Table S4). Within the target regions, 69% to 88% of cytosine positions were covered with the minimum depth of 3x, while 39%-79% were covered with the minimum depth of 10x, per sample (Figure S1). In respect to the transcriptomic data collected previously from the same samples (Banouh et al., 2023), the BS-seq data reported here covers 75–88% of differentially expressed genes (DEGs) with depths 3x or higher (Figure S2).

### Subgenome differences in mean methylation are inherited from lower-ploidy parents

Cytosine methylation occurs in the three sequence contexts (CpG, CHG and CHH) with different frequencies, as observed in other plant species. The highest mean TSS/SC methylation (considering only cytosines with ≥10x coverage) was observed at CpG sites (20%-27% across samples), with Tauschii-109 and Tauschii-87 being the most and least methylated, respectively. Methylation of the CHG sites ranges from 12% to 15%, while the methylation at CHH sites is the lowest, at 2%-3% (Figure S3). Since the mean methylation of the CHH context is close to the non-conversion rate (i.e. the proportion of unmethylated cytosines where bisulfite conversion failed), we ascertained the proportion of CHH sites where the methylation is statistically significant according to the binomial test. Across samples, the methylation level of 84%-90% CHH cytosines is not statistically different from the failed conversion rate, and only 10%-16% of all CHH sites around TSS/SC show significant levels of methylation (Table S5).

We calculated mean methylation levels per subgenome in all synthetic hexaploids, parental lines (Langdon and Tauschii-109), and the natural hexaploid wheat, using only cytosine positions consistently covered in all these samples (Figure 1; Figure S4). In all three sequence contexts, the B subgenome showed the highest mean methylation level, followed by A and D (Figure 1; Figure S4). This above-average B-methylation and below-average D-methylation is observed in both natural and synthetic hexaploids, as well as in the combined parental genomes. This suggests that the overall differences in subgenome methylation in the allopolyploids are not triggered by the whole-genome duplication, but rather are the legacy of the parental differences.

**Figure 1:**
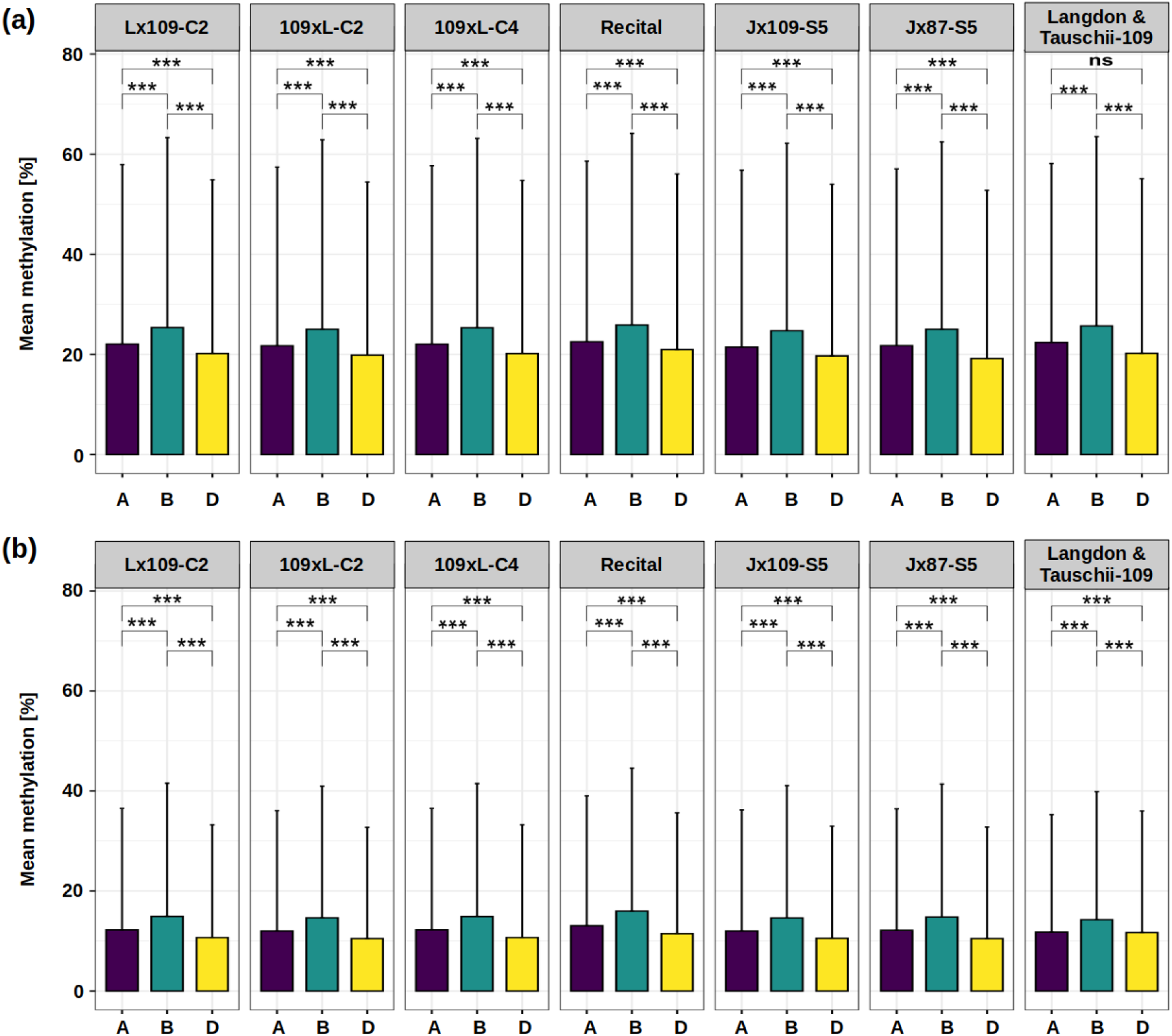
Mean methylation levels per subgenome (A, B and D) across genotypes in (a) CpG and (b) CHG contexts. Asterisks indicate significant differences (Wilcoxon test); ns, not significant at *alpha* = 0.05; *** *p* < 0.001. Error bars represent the standard deviation.

### TSS/SC methylation varies along chromosomes, but is stable across polyploidization

To examine DNA methylation landscapes along chromosomes in bread wheat, synthetic hexaploids and their lower-ploidy parents, we calculated weighted mean methylation across TSS and SC in 2Mb tiling windows and 1Mb sliding intervals, merging all sequence contexts into a single track. DNA methylation of the target regions varies considerably along chromosomes, with TSS/SC around centromeres showing elevated levels. Nonetheless, maximum values can sometimes be found away from centromeres (Figure 2a). Across samples, the chromosomes display highly similar methylation patterns, especially on A and B, with more apparent differences along the D chromosomes. Universally, differences between genotypes (Tauschii-87 vs. Tauschii-109; Recital vs. all other) are greater than differences between parents and their derived synthetics. The methylation landscapes of Langdon are highly correlated with the AB patterns of the derived synthetics (Pearson’s correlation in the range 0.992-0.995; Figure 2b). While the D methylomes of Tauschii-109-derived synthetics (109xL-C2, 109xL-C4, Lx109-C2, Jx109-S5) differ only slightly among each other (*r* range 0.991-0.995), all of them diverged more clearly from their D-parent (*r* range 0.963-0.978) (Figure 2c). This suggests that the D methylome encountered modifications that are consistent across multiple polyploidization events. Overall, the D methylome appears to be more diverse than the AB methylomes, and more prone to changes after polyploidization.

**Figure 2:**
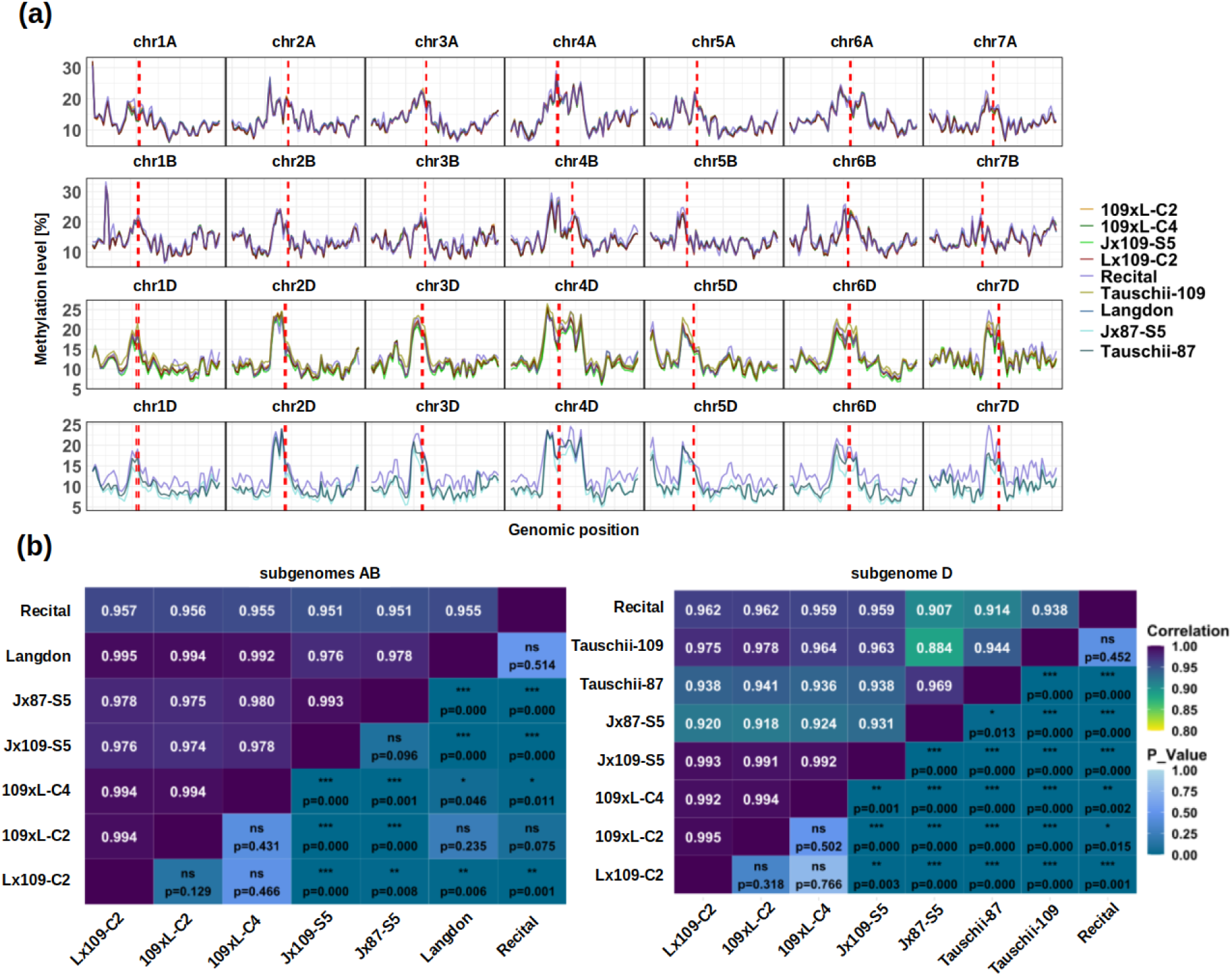
DNA methylation profiles and correlations across chromosomes and samples. (a) An overview of total DNA methylation along chromosomes (2Mb windows with 1Mb step), with each row of panels containing synthetic lines, their corresponding parental lines and the natural hexaploid Recital. The first and second rows display the A and B chromosomes, respectively, and both include the same synthetic lines derived from the AB parent Langdon as well as the corresponding AB chromosomes from Recital. The third row shows chromosomes from Tauschii-109, the D subgenome of the derived synthetics, and the D chromosomes of Recital. The fourth row represents chromosomes from Tauschii-87, the D subgenome of the derived synthetics, and the D chromosomes of Recital. (b) and (c) Heatmaps showing chromosome-level methylome differences between genotypes, separately for the D and AB subgenomes, respectively. Pearson’s correlation coefficients and Wilcoxon test p-values are given above and below the diagonal, respectively. Asterisks indicate significant differences: ns, not significant at *alpha* = 0.05; * p < 0.05; ** p < 0.01; *** p < 0.001.

### TSS/SC methylation does not clearly depend on centromere distance and is a poor predictor of transcriptional levels

In the following two sections, we investigated the TSS/SC methylation in relation to transcription and the distance from the centromere within trivial one-sample analyses (i.e. across all genes in a single sample, or jointly in all samples), not considering the polyploid context.

Since centromeric regions are rich in TEs and have relatively low gene densities, it can be expected that DNA methylation around centromeres is higher compared to more distal regions, as we have observed on the chromosome scans (Figure 2a). Accordingly, we observed a consistently significant negative correlation between CHG methylation and centromere distance for the group of genes with SC-centered data (Figure 3; Figure S5). However, this is not the case for CpG methylation around TSS, where we observed positive correlation (weak, but significant in all samples; Figure S5). In both cases, the proportion of shared variance is very low (*r^2^* <0.01), which means that TSS/SC methylation barely depends on the distance from the centromere. This contrasts somewhat with the relationship between gene expression [logarithmic form of transcripts per million (TPM)] and centromere distance, which is consistently negative and significant for both groups of genes (Figure S5). It may seem surprising that genes closer to the centromere have higher transcript levels, but this observation is consistent with findings of Zhao et al. (2023), explained by the high accessibility of centromeric chromatin.

**Figure 3:**
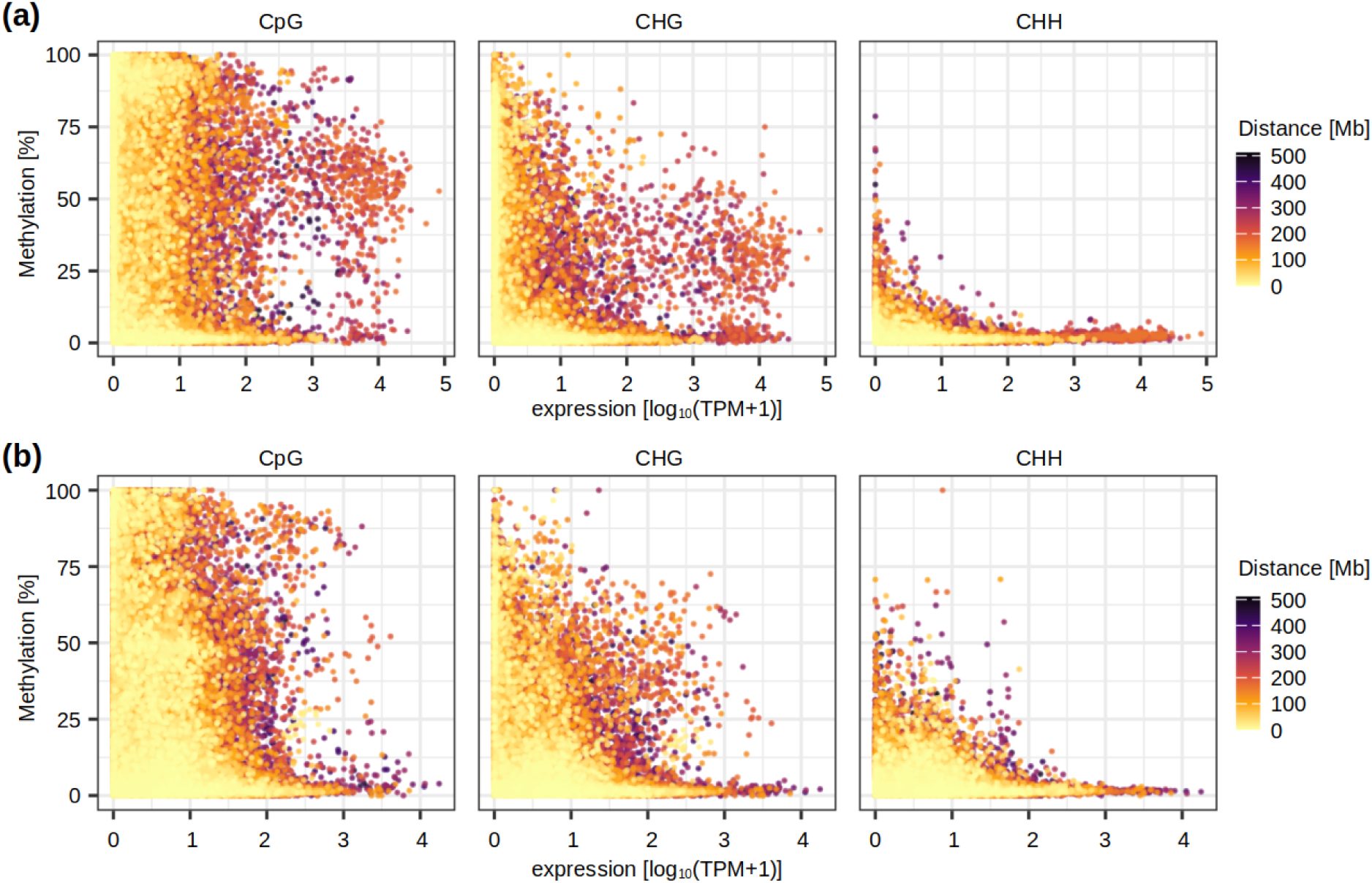
The relationship between transcription (x-axis), DNA methylation level (y-axis), and centromere distance (expressed with a color code). All samples with all three types of data were overlaid on the same scatter plot, and each dot represents a single gene. Data for individual samples, together with correlation coefficients for methylation-distance, methylation-expression, and expression-distance, are shown on Figure S5. (a) SC-centered dataset; (b) TSS-centered dataset.

The relationship between TSS/SC methylation and gene expression is consistently inverse and statistically significant in the CpG and CHG contexts of all samples (Figure 3; Figure S5). However, this relationship is very weak, with the proportion of shared variation (*r^2^*) between 0.004 and 0.025. For each sample, we observed large numbers of silent genes with both methylated and unmethylated TSS, as well as hundreds of moderately/highly expressed genes (TPM>10) with TSS methylation >25%. A group of genes with high expression (TPM >200) and medium-to-high methylation around SC (>30% in CpG and simultaneously >15% in CHG) can be recognized, with most of them (87) identified in at least two samples (Table S6). Among these genes, 34 are associated with the gene ontology term’nutrient reservoir activity’ (syn.’storage protein’; enrichment FDR 1.39E-85), which implies that genes producing seed storage proteins, which are highly up-regulated in the developing grain, have relatively high SC (i.e. gene body) methylation. CHH methylation shows only a weak relationship to transcription in the SC-centered dataset, and no consistent relationship in other comparisons (Figure 3; Figure S5).

### Fine-scale methylation around TSS/SC

Additionally, we explored the relationship between gene expression and DNA methylation on a finer spatial scale, summarizing DNA methylation in 10 bp intervals around TSS/SC. We separated expressed (TPM >0.1 in at least two libraries) and silent genes (TPM >0.1 in less than two libraries), creating four categories: (i) expressed genes with TSS-centered methylation data; (ii) silent genes with TSS-centered methylation data; (iii) expressed genes with SC-centered methylation data; (iv) silent genes with SC-centered methylation data. Subsequently, we averaged the 10 bp windows across all genes in a sample, and further averaged multiple samples to visualize a single pattern per category (Figure 4).

**Figure 4:**
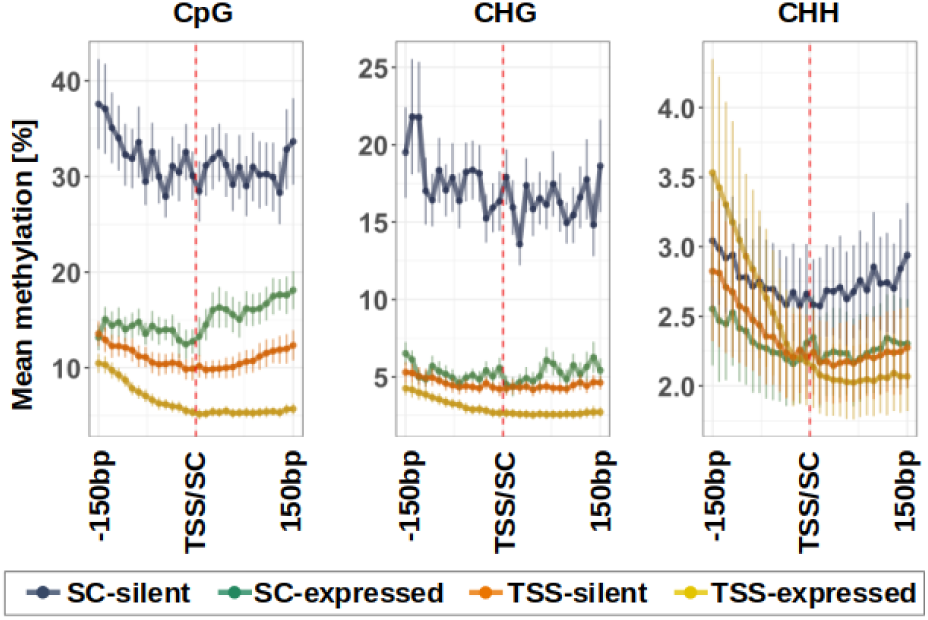
Mean methylation levels at ten-base pair resolutions around TSS and SC of all genes, averaged across all samples. The error bars represent standard deviations across samples (standard deviations across genes are much larger and difficult to visualize; see Figure S6).

Methylation levels around TSS are generally lower compared to SC. Within each dataset, expressed genes consistently show lower methylation levels than their silent counterparts (Figure 4). In general, DNA methylation levels are the lowest around TSS of expressed genes. Methylation progressively decreases as it approaches TSS and then remains low until 130 bp downstream. Compared to expressed genes, silent genes have their TSS methylation ∼5% and ∼2% higher in the CpG and CHG contexts, respectively. The methylation differences between expressed and silent genes are even more pronounced in the SC dataset, reaching ∼15% and ∼10% in the CpG and CHG contexts, respectively. This high difference is driven by a markedly higher SC methylation of silent genes, averaging around 30%-35% and 15%-20% in the CpG and CHG contexts, respectively. In other words, compared to TSS, local methylation patterns around SCs show a clearer methylation difference in relation to the gene expression status. While the pattern is consistent for the CpG and CHG context and the two groups of genes, the CHH context is more ambiguous, due to generally low methylation levels.

It is important to emphasize that Figure 4 shows double-averaged data sets (all genes are averaged per sample before averaging all samples), with standard deviations relating to the across-sample variation (not across-gene variation). Although the graphs appear to suggest that the expression category of individual genes can be confidently determined by checking the methylation status around TSS and SC, this is far from the truth, because across-gene variability in each of the four categories is much greater than the across-sample variability shown (Figure S6). Just as suggested by the scatter plots on Figure 3, each category contains genes with very high and very low DNA methylation.

### Homeolog expression bias is not caused by TSS/SC methylation

In the following three sections, we investigated TSS/SC methylation in the polyploid context, analyzing methylation and transcription of homeologs; recording methylation changes after polyploidization; and comparing differential gene expression and differential methylation between synthetics and their parents.

Triads and the homeolog expression contributions were used as described in Banouh et al., (2023), and compared with the weighted mean methylation level of TSS/SC regions of each gene. We considered a homeolog to have a’balanced’ contribution to the triad expression if its transcripts make up between 16.7% and 66.7% of the triad total. Below or above this range, the homeolog contribution is considered suppressed or dominant, respectively. When homeolog contributions are analyzed in the form of a continuous variable, very low and mostly insignificant correlation with DNA methylation is observed for all samples and sequence contexts (Figure 5a; Figure S7). Homeologs span the full range of methylation values for almost any level of expression contribution. It appears that with increasing dominance, homeologs are associated with lower methylation levels. However, it should be noted that very few homeologs show strong dominance, therefore the apparent pattern can be due to decreasing number of data points. The lack of correlation suggests that methylation levels do not determine the level of contribution to the overall triad expression.

**Figure 5:**
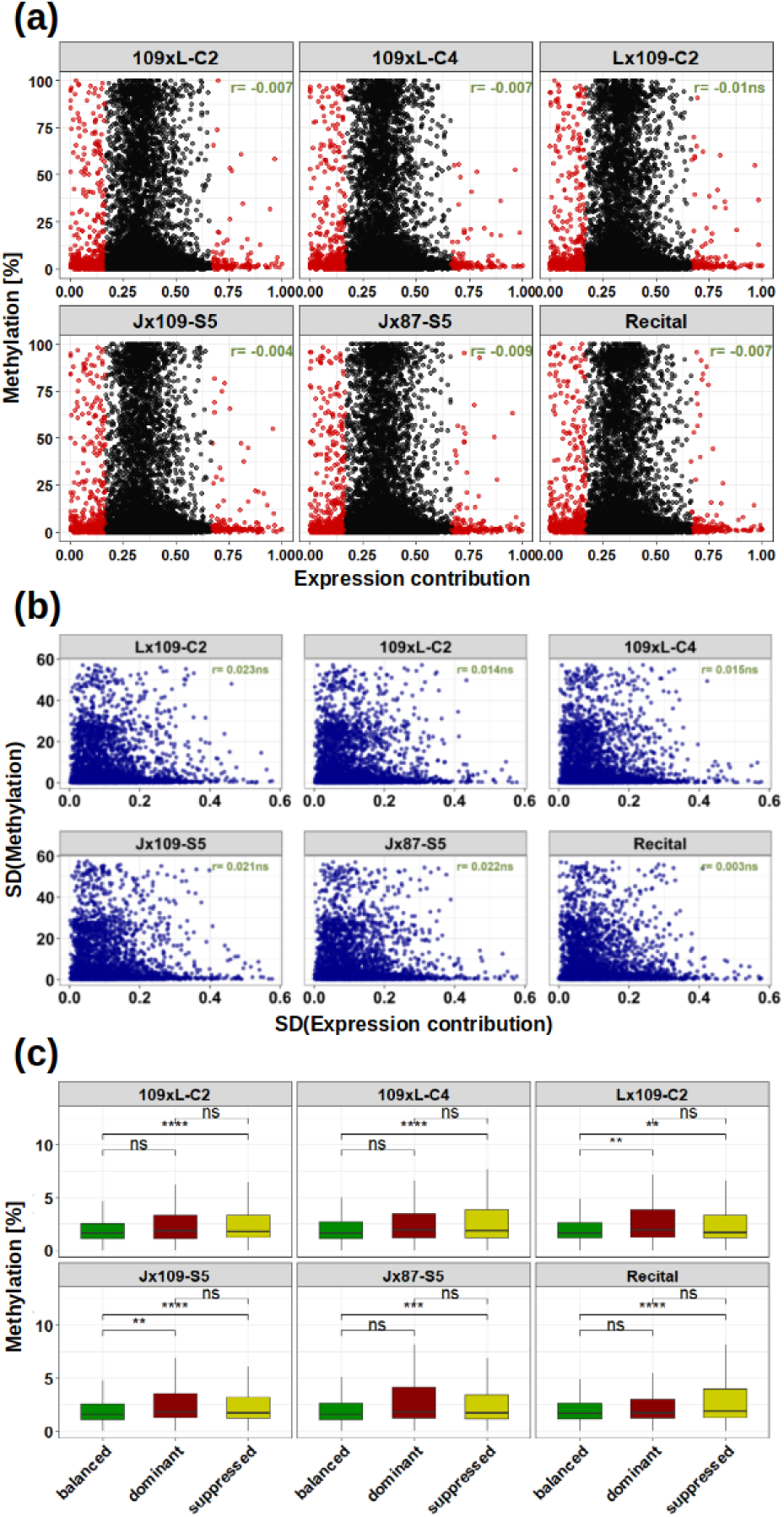
DNA methylation and homeolog expression bias. (a) Scatter plots with each data point representing a single homeolog, with its contribution to triad expression on the x-axis and CpG methylation level on the y-axis. Suppressed and dominant homeologs are shown in red. (b) Scatter plots with each data point representing a triad, with variance (SD) in the expression contribution on the x-axis, and variance of the CpG methylation level on the y-axis. Correlation coefficients are shown in green. (c) Boxplots showing CpG methylation across balanced, suppressed and dominant genes. Asterisks indicate statistical significance (Wilcoxon test; ** *p* < 0.01; *** *p* < 0.001; ns, not significant at *alpha* = 0.05).

Since the scatter plots with individual homeologs dissolve the triad information (i.e. homeologous groups are not identifiable on the scatter plot), we further checked whether the variance of methylation within triads correlates with the variance of transcription. Such a comparison tests the hypothesis that imbalanced triads are associated with high methylation differences while uniform triad methylation leads to a balanced homeolog expression, and is therefore independent from the direction of the relationship (i.e. it does not matter whether high methylation causes low transcription in one triad, but high transcription in another triad). This comparison did not detect significant correlations in the CpG and CHH contexts, and found significant but very weak positive correlations in the CHG context (proportion of shared variation *r^2^* ∼0.003) (Figure 5b; Figure S8). This does not suggest that DNA methylation plays an important role in homeolog expression differences. The relationship is not clearer when homeolog contributions are analyzed in the form of categorical data, creating balanced, dominant and suppressed homeolog groups (Figure 5c). While the mean CpG methylation of suppressed homeologs is consistently distinguished from the mean of balanced homeologs (*t*-test; alpha = 0.05), dominant and suppressed homeologs are not statistically different in any of the allohexaploids. This pattern is repeated in the CHG and CHH contexts (Figure S9). The result showing that suppressed homeologs are statistically different from balanced homeologs but not from the dominant ones is probably related to differences in group sizes, but nonetheless perplexing, as the dominant and suppressed categories are expected to be the most distinct.

### DNA methylation changes in nascent polyploids

We searched for DMRs (differentially methylated regions) between synthetics and their parents, analyzing each cytosine context separately, with two different approaches defining the’methylation regions’. The first approach uses the entire target regions (300 bp around TSS/SC) as fixed-length windows. The second approach is based on a change point segmentation calculated with the methseg function in MethylKit. It determines segments of varying sizes defined by the homogeneity of methylation levels across consecutive cytosines. The segments were identified from pooled parental data, and were set to contain a minimum of three CpG sites with ≥10x coverage. These segments were then used to track methylation changes across the polyploidization event.

The change point segmentation approach identified between 34 and 47 DMRs within sets of 44,944-45,835 genes analyzed (Figure 6a). The CpG context recorded the highest number of TSS/SC DMRs, with counts ranging from 21 to 32 per sample, followed by the CHG (8-15) and the CHH context (1-2). Across samples, 109xL-C2 recorded the highest number of DMRs (1-32 depending on the context). In respect to subgenomes, most changes occurred on D, with 17-26 DMRs across samples. This is in agreement with the comparison of general methylation patterns along chromosomes (Figure 3). The ratios of hypermethylation and hypomethylation were not consistent across the different synthetics. In Lx109-C2, hypomethylation was more common than hypermethylation for all subgenomes. In contrast, hypomethylation was more prevalent in 109xL-C2, but this pattern did not persist in the subsequent generation (109xL-C4). Some of the identified DMRs occur repeatedly in multiple synthetics (involving 3-7 genes per pairwise comparison; Figure 6b). Such overlaps are statistically significant according to the hypergeometric test, which predicts <2 genes to overlap by chance in sets of the given sizes. This indicates that some nonrandom and reproducible TSS/SC methylation changes occur in the synthetics. Nonetheless, it should be noted that significant methylation changes in parent-synthetic comparisons were detected in a very small fraction of the analyzed genes (0.08%-0.1%).

**Figure 6:**
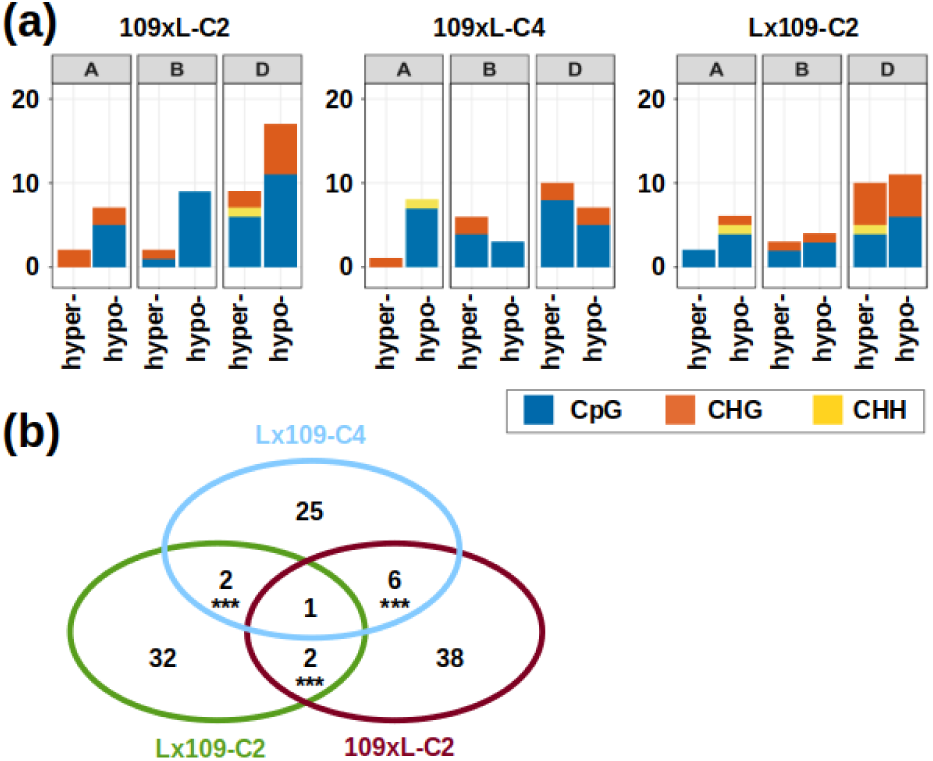
DMR identification using the change point segmentation approach. (a) Counts of hyper/hypomethylated DMRs in different hexaploid synthetics across different subgenomes and different cytosine contexts. (b) Venn diagram showing the overlap of DMRs (considering all contexts) detected in different synthetic samples. Asterisks indicate the significance of the overlaps (*** *p* < 0.001).

When the entire target windows of 300 bp around TSS/SC were used as regions in the DMR analysis, slightly more DMRs were identified (91-110; Figure S10a), corresponding to 0.09%-0.1% of the total genes analyzed. While the patterns seen in the change point segmentation were preserved here (i.e. the excess of changes on the D subgenome), the two methods detected generally different sets of genes (Figure S11). However, it is difficult to assess the sensitivity and specificity (the false negative and false positive rates, respectively) of DMR detection methods, since it is unclear what a’real DMR’ (i.e. true positive) is. A real DMR could be defined as a DMR with biological consequences (e.g. transcriptional change, chromatin structure,…); however, we cannot assess biological consequences of DMRs prior to their identification, which introduces a circular problem. For the moment, any method of DMR detection that identifies statistically significant differences may be considered valid, but we have to accept the possibility that some or all of the detected DMRs may have no biological consequences.

### DEGs and differentially methylated TSS/SCs are unrelated in nascent wheat polyploids

In our previous study (Banouh et al., 2023), we identified sets of DEGs triggered by polyploidization in wheat synthetics. With DNA from the same nucleic acid extracts that had been used for the transcriptomic analysis, here we searched for DNA methylation changes and tested whether the changes in transcription can be explained by mC modifications. Only DEGs with sufficient coverage in the BSseq analysis (10x) were included in the comparison. The association between DEGs and DMRs was assessed with Venn diagrams (Figure 7a) and the exact hypergeometric test. Across all samples and either of the DMR detection methods, only four genes simultaneously change their methylation and transcription after polyploidization: TraesCS2D02G309500 (12-oxophytodienoate reductase2) downregulated with hypermethylated TSS; TraesCS5D02G553000 (beta-fructofuranosidase) downregulated with hypomethylated SC; TraesCS6D02G026500 (acid beta-fructofuranosidase) downregulated with hypermethylated TSS (all three in Lx109-C2); and TraesCS3B02G234600 (Vesicle transport v-SNARE 13) downregulated with hypermethylated SC (in 109×L-C4). However, it is unclear whether the differential expression and methylation in these genes are causally related. According to the hypergeometric test, such overlaps are not significantly different from random, given the sizes of the DMR and DEG subsets. Therefore, we cannot reject the null hypothesis regarding the independence of DEGs and DMRs.

**Figure 7:**
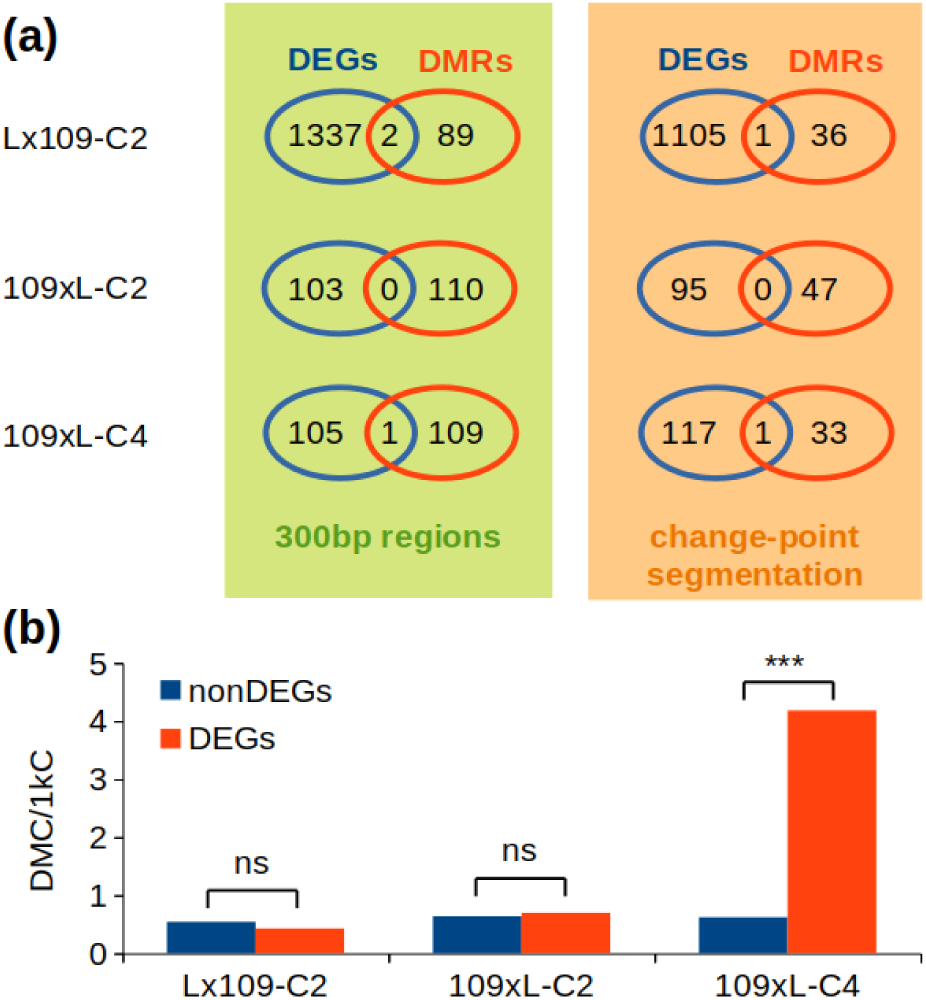
Differential expression and differential methylation. (a) Venn diagrams representing per-sample overlaps between DEGs and DMRs obtained with either of the two methods used. Numbers of DEGs differ between the two DMR detection methods because only DEGs with sufficient DNA methylation data are included in the statistical comparison. None of the overlaps is statistically over-represented according to the exact hypergeometric test with normal approximation. (b) Occurrence of DMCs in DEGs and non-DEGs, expressed as the proportion of DMCs within 1,000 covered cytosines. Statistical differences between DEGs and non-DEGs were tested with the hypergeometric test and the chi-square test; ns, not significant at *alpha* = 0.05; *** *p* <0.001.

In addition to the DMR analysis, we also investigated the proportion of DMCs (differentially methylated cytosines) in DEGs and non-DEGs. A cytosine with ≥10x coverage was considered a DMC if the methylation level difference between parents and synthetics was ≥ 25% and *p* < 0.01. We detected 0.43 and 0.7 DMCs per 1,000 DEG-associated cytosines in Lx109-C2 and 109xL-C2, respectively, which is not statistically different from the frequency of DMCs in non-DEGs (Figure 7b). In 109xL-C4, we found that the frequency of DMCs was significantly higher in DEGs (compared to non-DEGs; chi-square test and hypergeometric test). This was mainly due to 19 hypermethylated DMCs around the SC of TraesCS3B02G234600 (Vesicle transport v-SNARE 13). This gene is downregulated in 109xL-C4, where it was also detected as DMR (300bp regions method). It is therefore possible that the hypermethylation and downregulation are causally related in this case. Overall, our results show that methylation changes around TSS/SC are rarely induced by allohexaploidization in AABBDD-type wheat synthetics, and perhaps with a single exception are not statistically associated with changes in gene expression.

## DISCUSSION

### The unclear role of DNA methylation in gene expression and polyploid formation

DNA methylation is regarded as one of the crucial mechanisms of epigenetic regulation of gene expression. In mammals, hypermethylation of CpG islands in promoters can cause transcriptional silencing, which is in particular cases associated with cancer development (Das and Singal, 2004, Bernstein et al., 2013). Hypermethylation is also involved in the formation of facultative heterochromatin, best exemplified by the X-chromosome inactivation, where DNA methylation is employed to maintain (though not to initiate) stable silencing that compensates the gene dosage imbalance in mammalian females (Chow et al., 2005). Due to its evolutionary conservation and taxonomic ubiquity, DNA methylation is assumed to play similar roles in plants. Nonetheless, particular features of plant DNA methylation, namely the occurrence of mC in any sequence context (symmetrical CpG and CHG, or asymmetrical CHH) and the absence of CpG islands, make the study of DNA methylation in plants more difficult, and its role in epigenetic regulation more complex.

DNA methylation has long been hypothesized to play an important role in polyploidization, which is a frequent phenomenon in the plant kingdom. Although polyploidization does not create a gene dosage imbalance, as copy numbers of all genes are multiplied equally, it is often assumed that nascent polyploids need to fine-tune their gene expression in the new genomic environment, and that epigenetic changes are involved in this process. It has been speculated (Matzke et al., 2015) that the RNA-directed DNA methylation (RdDM) pathway could be involved in the’diploidization’ of polyploid genomes, i.e. a gradual evolutionary process of sub-/neo-fuctionalization and loss of redundant gene copies. Conceivably, RdDM triggered by the’shock of genome doubling’ could selectively inactivate redundant gene copies, leading to heterochromatin formation, and subsequently to sequence diversification or gene loss (Matzke et al., 2015). Following similar hypotheses, DNA methylation patterns have been investigated in a number of plant species with whole-genome doubling in their ancient history, including rice (Wang et al., 2013), maize (Zhao et al., 2017; Renny-Byfield et al., 2017), soybean (Zhao et al., 2017), pear (Li et al., 2019b), and *Brassicaceae* (Parkin et al., 2014; Chalhoub et al., 2014), usually finding little evidence of epigenetic differentiation of subgenomes, and an unclear relationship between DNA methylation and transcriptional activity of the duplicated genes (Bellec et al., 2023).

The study of methylomic changes after polyploidization presupposes a more general understanding of the role of DNA methylation in plants. The impact of cytosine methylation on gene expression is of particular interest, but the relationship is still shrouded by uncertainty. Given the absence of distinct CpG islands in plant promoters and plant genomes in general, there is typically no prior information on the position of elements whose methylation has consequences for the transcription of adjacent genes. DNA methylation is therefore typically measured in the transcribed gene partitions (gene bodies) and loosely-defined promoters. However, gene body methylation (gbM) is persistently difficult to relate to transcription, not only in the polyploid context (Parkin et al., 2014; Chalhoub et al., 2014; Zhao et al., 2017), but also in diploids. After detailed analyses of gene expression and methylomes in *Arabidopsis thaliana* epigenetic recombinant inbred lines and in *Eutrema salsugineum* that lacks gbM, Bewick et al., (2016) found no evidence supporting gbM involvement in the regulation of transcription. Nonetheless, gbM could be involved in the prevention of aberrant and spurious transcripts that initiate outside of proper TSS, as it is described in mammals (Neri et al., 2017; Wu et al., 2022). This idea is consistent with the hypothesis that the initiation of transcription requires unmethylated DNA and is prevented in methylated regions. This is in agreement with observations of low methylation around TSS across multiple plant species (Niederhuth et al., 2016), but this general pattern does not indicate whether TSS methylation can be employed by plants as an epigenetic on-off switch to fine-tune gene expression under internal (genomic) or external (environmental) stress.

### TSS methylation is only weakly associated with gene silencing

We hypothesized that the region around TSS, which is the place of transcriptional initiation and includes the’core promoter’, could be crucial for the epigenetic control of gene expression. Focusing on 300 bp windows centered around TSS, we observed a wide range of CpG methylation for virtually any gene expression level (Figure 3; Figure S5). Although most wheat genes have mean CpG methylation of their TSS below 10%, there are typically >1,000 outliers with TSS methylation 25-100% and transcript abundance in the range 1×10^0^-1×10^4^ TPM. Although CpG methylation and gene expression are negatively correlated with statistical significance in all samples, the proportion of shared variation (*r^2^*) is extremely low (0.009-0.021) and hardly indicative of a causal relationship. The situation is analogical in the CHG context, and the scatter plots of transcript abundance and DNA methylation around SC shows a similar distribution. In fact, gene expression is as hard to relate to DNA methylation around TSS/SC, as it is to gene body methylation reported in other species (Parkin et al., 2014; Liang et al., 2019). On a finer spatial resolution (Figure 4), across-gene means of CpG and CHG methylation at TSS can distinguish expressed (TPM >0.1) and silent states, but the difference in methylation is even more pronounced at the SC, with very high variance in both cases.

These results seem to be in contrast to those reported in Niederhuth et al., (2016), who quantified for multiple species transcript abundance in mCG-TSS genes, i.e. genes where TSS overlaps with mCG regions (defined on the basis of a binomial test). For most examined species including *Brachypodium distachyon* and *Zea mays*, transcript abundance of mCG-TSS genes is virtually zero, although it should be noted that the boxplots (Figure S19 in Niederhuth et al., 2016) do not show outlier data points. Our results indicate that wheat is probably more similar to *Oryza sativa* or *Solanum lycopersicum*, where CpG methylation of TSS permits moderate levels of transcription. The poor correlation between TSS methylation and gene expression in diploid, tetraploid and hexaploid wheats observed here, and the variability of this relationship across genera do not suggest that TSS methylation has universally strong consequences for the level of gene expression.

This conclusion adds further ambiguity to the presumed relationship between transcription and DNA methylation. The absence of a clear genome-wide methylome-transcriptome pattern are surprising, given some *in vitro* studies (O’Malley et al., 2016) and the evolutionary conservation of DNA methylation pathways. Below, we consider several explanations for this persistently obscure relationship, possibly working in combination.

i. DNA methylation controls the expression in only a fraction of genes and has no impact in most cases. There is strong evidence to support this possibility. In human cells, methylation of only 16.6% of CpG sites shows a significant correlation with the transcription at neighboring TSS (Medvedeva et al., 2014). The authors concluded that’*direct and selective methylation of certain TF binding sites that prevents TF binding is restricted to special cases and cannot be considered as a general regulatory mechanism of transcription*’. In plants, the global effect of DNA methylation on transcription can be assessed with mutant lines of *A. thaliana* defective in DNA methylation maintenance. In DNA methylation-free quintuple mutant with all five known DNA methyltransferases (MET1, DRM1, DRM2, CMT3, and CMT2) knocked out, extreme growth retardation and small size was observed, together with a suite of severe developmental defects (He et al., 2022). Nonetheless, the complete removal of DNA methylation caused only 3,738 DEGs (fold change >2; FDR <0.01), which is 13.6% of all annotated protein-coding genes. Much fewer DEGs (310; 1.1% of all annotated protein-coding genes) were observed in the quadruple mutant (*drm1 drm2 cmt3 cmt2*) retaining CpG methylation but defective in CHG and CHH methylation maintenance (He et al., 2022). Since the complete removal of DNA methylation modifies the transcriptional regulation of only a fraction of genes, local and lower-magnitude changes of DNA methylation are likely to have even smaller impacts on transcription.
ii. The effect of DNA methylation on transcription levels is mild, frequently escaping detection in low-powered studies. It is possible that DNA methylation affects transcription on a very fine scale, with meaningful impacts only for lowly-expressed genes. Such small changes could be undetectable in studies with low sensitivity due to a small number of biological replicates, or the signal could be overwhelmed by highly-expressed genes where DNA methylation has no substantial effect.
iii. The role of DNA methylation in the regulation of gene expression is complex, masking clear genome-wide patterns. The complexity can be manyfold. DNA methylation of a regulatory element can prevent the binding of both transcriptional activators and repressors, potentially leading to silencing and upregulation, respectively (Zhang and Zhu 2024). Moreover, some TFs show affinity to methylated binding sites (O’Malley aet al., 2016). Consequently, high DNA methylation can be conducive to high levels of transcription in some genes, but low levels in others, and the same can be true for low DNA methylation. This is consistent with the DEG analysis of the Arabidopsis quintuple mutant, where demethylation causes nearly as many down-regulations as up-regulations (He et al., 2022). However, the down-regulations (and up-regulations alike) could be an indirect result of the demethylation. In the subsequent study of the quintuple mutant, Zhao et al., (2022) demonstrated that the complete loss of DNA methylation causes extensive switches of chromatin states. The differential expression of thousands of genes may then result from chromatin switches or increase/decrease of histone modifications, rather than solely from DNA demethylation. When histone modifications act independently from mC, the link between DNA methylation and transcription is further eroded.
iv. DNA methylation drives transcriptional changes dynamically in response to challenge (genomic or environmental stress), and the effect is invisible in a static sample. In this scenario, basic gene expression levels are generally determined by non-epigenetic regulatory networks, but are responsive to DNA methylation changes under stress conditions. This type of association can be explored by comparing sets of DEGs and DMRs in multiple-sample comparisons, with examples involving various stresses and plant species (Xu et al., 2017; Feng et al., 2016; Wang et al., 2016). A methylation-driven stress response can be concluded from a significantly over-represented DEG-DMR group (using a 2×2 chi-square contingency table, or the hypergeometric test); however, the significance is rarely tested in published studies. Here, we explored the question whether transcriptomic changes induced by polyploidization (presumably causing genomic stress) are associated with TSS methylation changes.

### Changes in gene expression and TSS/SC methylation are independent in nascent wheat allohexaploids

Polyploidization in wheat triggers very few DNA methylation changes around TSS and SC. When comparing synthetics (including reciprocal crosses and two different generations) to their parents, we detected only 91-110 DMRs among ∼93,000 genes with sufficient methylation data (considering 300 bp TSS/SC windows as regions). Most of the DMRs and DMCs were consistently found on the D subgenome, which is similar to the distribution of DEGs detected in the same samples (Banouh et al., 2023). This indicates that the D subgenome of nascent wheat allohexaploids is more vulnerable to both transcriptomic and methylomic changes. However, the DMRs and DEGs are not statistically related, since we found only 0-2 genes per synthetic where differential expression is accompanied by differential methylation.

### Homeolog expression differences in wheat are not driven by DNA methylation

Among the three subgenomes of natural and synthetic allohexaploid wheat, B and D are the most and the least methylated, respectively, with statistical significance. It is tempting to relate this observation to the subtle yet significant D-dominance in transcript abundance observed in the same samples (Banouh et al., 2023), as well as the Chinese Spring reference genome (Ramirez-Gonzales et al., 2018). Moreover, when assessing homeolog expression bias in Banouh et al., (2023), we observed a statistically strong under-representation of D-suppressed triads, sometimes accompanied by an excess of B-suppressed triads. We have demonstrated that homeolog expression imbalance is mainly due to parental legacy, i.e. it is not a consequence of polyploidization but merely a perpetuation of pre-existing parental expression differences. Similarly, here we demonstrate that the subgenome methylation differences are not a consequence of polyploidization, since the A, B and D subgenomes are also methylated at different levels in the parental genotypes. It is interesting that the methylomes of the highly syntenic subgenomes were not homogenized after ∼10,000 years of coexistence in natural bread wheat, or even after 0.5 million years in the case of the A and B methylome differences in tetraploid wheat. It indicates a very faithful long-term maintenance of DNA methylation patterns. This observation, together with the below-average D-methylation, the subtle D-dominance and the dearth of D-suppressed triads suggest that homeolog expression bias could be driven by DNA methylation differences.

However, here we show that the relationship between DNA methylation and homeolog contribution to triad expression is either absent, or only weakly significant (Figure 5a; Figure S7). When the homeolog contributions are regarded as a qualitative variable, the dominant and suppressed copies are statistically indistinguishable on the basis of TSS/SC methylation in most cases (Figure 5c; Figure S9). Considering the duality of the possible effects of DNA methylation on transcription (leading to up-and down-regulation in different cases), we also checked the correlation between methylation and expression variance within triads (Figure 5b; Figure S8). This comparison does not assume a dominant effect of methylation (be it gene activation or silencing), it merely checks whether triads with unequal homeolog contributions are also characterized with high variability of DNA methylation. This is certainly not the case for CpG methylation nor for CHH methylation. We detected significant, but very low positive correlation in the CHG (proportion of shared variance *r^2^*∼0.003), not indicative of a causal relationship.

Our conclusion that homeolog expression bias in bread wheat and analogical synthetics is not driven by DNA methylation is consistent with observations in more ancient polyploids. Zhao et al., (2017) studied homeologs in the soybean genome that went through a polyploidization 5-13 million years (Mya) ago, and found that gene pairs with >2 fold expression differences are not significantly different in their patterns of DNA methylation, or the distribution and abundance of 24-nucleotide siRNAs. Furthermore, Parkin et al., (2013) found that the patterns of cytosine methylation in paralogs on different subgenomes of *Brassica oleracea* (hexaploidization 23 Mya) do not correlate with individual gene dominance. In a more recent polyploid *Brassica napus* (allotetraploidization 7,500 years ago), homeologs with differential UTR or gene body methylation are equally expressed in over half of the cases (Chalhoub et al., 2014). Although independence of homeolog expression and methylation in the CpG and CHG context is statistically rejected for gbM, with highly-methylated/lowly-expressed and lowly-methylated/highly-expressed homeologs in excess, the violation of independence is much weaker for UTR methylation (this was not shown in the original study, but a chi-square test can be performed on data in Table S45 of Chalhoub et al., 2014). In the maize genome (polyploidization ∼10 Mya), the more highly expressed gene in the homeolog pair typically exhibits a lower proportion of mCs up-stream of TSS and in gene bodies, and a more open chromatin (Renny-Byfield et al., 2017), though the authors emphasize that the attribution of cause and effect was impossible. On the other hand, a causal link between gbM and expression of homeologs is claimed by Wang et al., (2017). The authors studied a link between gbM divergence and expression difference of paralogs in rice (polyploidization 70 Mya), using *OsMet1-2* null mutant defective in CpG methylation maintenance. They found that duplicated gene copies correlate with each other in their expression levels more strongly once CpG methylation is removed (a change from *r* = 0.43 to *r* = 0.51). However, while Wang et al., (2017) focus on gbM, which they claim to be the cause of the differences in duplicate gene expression, CpG methylation in the *OsMet1-2* null mutant is lost globally, not just in gene bodies. It is therefore unclear whether the observed reduction in expression differences (which is rather small anyway) is connected to demethylation of gene bodies, regulatory elements, TEs, or even to changes in chromatin states that are triggered by global demethylation (as shown by Zhao et al., 2022).

Our results and the overview of different polyploid genomes suggest that the association of homeolog expression differences with DNA methylation is sometimes detectable on a statistical basis, but is so weak that the transcriptional divergence of most homeologs cannot be attributed to DNA methylation differences. What then is the main cause of homeolog expression bias is a curious question. Homeologs, especially those in allohexaploid wheat, have high sequence similarity, which suggests that they share the same regulatory networks in most cases. It is therefore unlikely that homeolog expression bias is a consequence of different regulatory mechanisms (a TF regulating a particular gene is very likely to have the same effect on all homeologous copies). Thus, whether the frequent homeolog expression differences are driven by chromatin states, 3D-positioning of chromatin within the nucleus, or other unknown factors, remains open for future research.

### Wheat genome unfazed by polyploidization

Polyploidizations are commonly believed to cause a’genome shock’, i.e. a widespread disturbance of gene regulation and genome integrity related to changes in gene dosage or parental incompatibilities (McClintock 1984; Tayalé and Parisod 2013). Accordingly, allopolyploidization in wheat has been associated with rampant transcriptomic changes (Zhang et al., 2016; Hao et al., 2017; Vasudevan et al., 2023), sequence loss (Ozkan et al., 2001; Ozkan et al., 2003), activation of TEs (Kashkush et al., 2002) and epigenetic changes (Yaakov and Kashkush 2010; Kraitshtein et al., 2010). However, a genome-wide analysis of TEs in the bread wheat genome found no signs of transpositional bursts after allohexaploidization (Papon et al., 2023), and we have previously reported that only about 1% of genes significantly change transcription in nascent AABBDD allohexaploids, without a statistical relationship to adjacent TEs (Banouh et al., 2023). Here we reported that DNA methylation differences around TSS and SC are extremely rare (∼0.1% of genes) in comparisons of early generations of wheat allohexaploids and their parents. These observations do not support the notion of a genome shock in polyploid wheat, and we consider several reasons why our results contradict earlier studies.

i. First of all, large over-estimations of transcriptomic response to polyploidization are often caused by technical difficulties associated with interploidy comparisons (see Supplementary Note in Banouh et al., 2023).
ii. Secondly, studies conducted prior to the assembly of the wheat genome provided only a partial view of the changes, without a genome-wide perspective. A hunt for changes overlooks the fact that the vast majority of the genome remains unchanged.
iii. Finally, it is very likely that the magnitude of changes depends on parental genotypes. Since there is some evidence that allopolyploidizations trigger much stronger genomic responses than autopolyploidizations (Tayalé and Parisod 2013), more diverged parental combinations can be expected to cause more changes than the less diverged ones. It may be possible to find a particular combination of *Ae. tauschii* and tetraploid wheat (e.g. *T. dicoccoides*) that leads to a massive genomic reorganization, or one that is entirely incompatible. However, if the goal is to find out what actually happened in the genomic history of bread wheat, there is no reason to assume such parental incompatibilities. In our assessment, changes caused by allohexaploidization in wheat may not be qualitatively or quantitatively different from those triggered by intra-specific hybridizations.

## MATERIALS AND METHODS

### Plant material

All seeds used in this study were provided by the ‘Centre de Ressources Biologiques Céréales à Paille’ (Small Grain Cereals Biological Resources Centre), INRAE, Clermont-Ferrand, France. All data reported here were obtained from the developing grain sampled at 250 degree-days from both natural and synthetic hexaploid wheat lines (AABBDD) as well as from the parental genotypes *Ae. tauschii* (DD) and *T. turgidum* subsp. *durum* (AABB) (Table S1). Natural hexaploids are represented by the bread wheat cultivar’Recital’. The synthetic wheats were prepared by crossing *Ae. tauschii* genotype’Tauschii-109’ with the durum cultivar’Langdon’ (producing synthetics’109xL-C2’,’109xL-C4’ and’Lx109-C2’, with identifiers indicating the direction of the coss and the sampled generation after colchicine-induced chromosome doubling). Additional spontaneous allohexaploids’Jx87-S5’ and’Jx109-S5’ produced by a different research group (INRAE - Agrocampus Rennes) from the durum cultivar’Joyau’ and *Ae. tauschii* genotypes’Tauschii-109’ and’Tauschii-87’ were also analyzed, although direct descendants of the parental lines used for the crosses were not available for this study. Additional details about the allohexaploid synthesis, nomenclature and growth conditions are given in our previous paper Banouh et al., (2023) reporting on the transcriptomic analysis of these samples.

### DNA extraction

The nucleic acid extracts that had been used previously in Banouh et al (2023) for RNA purification were processed here to obtain genomic DNA. Briefly, the grain tissue was ground into a fine powder using liquid nitrogen, a mortar and a pestle. Up to 1 g of the powder was dissolved in 4.5 ml of extraction buffer (0.1 M NaCl, 10 mM Tris-HCl, 1 mM EDTA, 1% SDS). Nucleic acids were extracted twice with phenol:chloroform:isoamylalcohol (25:24:1) and subsequently precipitated using 3 M sodium acetate (1/10 volume) and two volumes of absolute ethanol. Following centrifugation, the resulting pellet was resuspended in nuclease-free water. An aliquot of this crude nucleic acid extract (50–75 µg) was used for the RNA-seq purification (Banouh et al., 2023), while another one was treated with 30 µg RNase A and subsequently purified with silica-based spin columns (Qiagen) in order to obtain purified DNA. DNA was obtained from two biological replicates per sample.

### Capture-BSseq probe design

Transcription start sites were identified on the basis of the Chinese Spring v1.1 gene annotation, treating the first position of 5’ untranslated regions (5’-UTR) as TSS. Out of 107,891 annotated HC genes, 68,731 (63.7%) have 5’-UTR annotations. In the remaining 36.3% of HC genes, TSS information is missing due to the absence of full-length mRNA data, and we aimed to capture the beginning of the coding sequence instead, i.e. the SC of those genes. While the SC position is not relevant for the initiation of transcription, the SC regions provide an insight into 5’-UTRs (upstream of SC) and the coding sequence (downstream of SC). Each gene was targeted by two probes within a 300 bp window around TSS or SC (one probe upstream and another one downstream). In order to increase the applicability of the probe design to varied *Triticeae* genomes (diploid *Ae. tauschii*, tetraploid *T. turgidum* sp., hexaploid *T. aestivum*), the probe design was adjusted according to 15 different chromosome-level assemblies - Chinese Spring v1.1 (IWGSC 2018); 10+ Wheat Genome Project (Walkowiak et al., 2020); *T. turgidum* subsp. *durum* cv. Svevo v2 (Maccaferri et al., 2019); *T. turgidum* subsp. *dicoccoides* (Avni et al., 2017); and two assemblies of *Ae. tauschii* v4.0 (Luo et al., 2017) and AOCO02000000 (Zhao et al., 2017). Our design aimed for probes that correspond to consensus TSS/SC regions of all these assemblies rather than just the Chinese Spring reference. To achieve this, all assemblies were shredded into 500 bp fragments with 250 bp sliding step using GenomeTools shredder, and all fragments were mapped onto the Chinese Spring v1.1 reference with bwa mem (Li, 2013), creating a single bam file. Reads with mapping quality <20 were filtered out (samtools). Subsequently, fragments (or rather a single consensus sequence per genome built from overlapping fragments) around each TSS/SC were extracted from the bam file in the form of multiple sequence alignments, using a custom python script (150 bp upstream and 150 bp downstream). All upstream fragments (for all genes and genomes) were then re-assembled into contigs with Geneious 6.1 (Biomatters; http://geneious.com) allowing no gaps and up to 3% mismatches within ≥120 bp overlaps. The same procedure was repeated for all downstream fragments. These contigs were considered as probe candidates, and further filtered. We removed probe candidates (i) redundant among the upstream and downstream sets; (ii) mapping to the wheat chloroplast genome with ≤10% mismatches; (iii) having valid alignments (bwa mem) with <8 of the used assemblies; and (iv) having more off-target than on-target blat hits (using-fastMap and - minScore=80). This process resulted in 200,495 consensus probes covering ∼100k genes of wheat (typically with 2 probes per TSS/SC). Chinese Spring genes corresponding to all the removed probe candidates were not enriched in any GO terms (using TGT website http://wheat.cau.edu.cn/TGT/; Chen et al., 2020), indicating that our probe design is unbiased in respect to gene function. The capture probes were synthesized by Agilent (SureSelect custom design), covering both strands of the targets.

### Preparation, sequencing and mapping of the capture-BSseq libraries

The sequence capture protocol with SureSelect Custom Tier6 probes (Agilent) was applied on genomic DNA prior to the bisulfite conversion with EZ DNA Methylation-Gold Kit (ZYMO Research). Illumina-compatible libraries were prepared with SureSelectXT Methyl Reagent kit (Agilent), with post-conversion multiplexing for parallel sequencing of all samples. The libraries were prepared by PGTB (France) and sequenced on NovaSeq6000 (2×150) by Genotoul (France) to a theoretical depth of target coverage 40x. We obtained raw reads from 18 directional libraries (with both the top and bottom DNA strands). These reads underwent initial processing with Trim Galore to eliminate adapter sequences, reads with length <20, and unpaired reads. Reads passing these quality filters were aligned to the Chinese spring reference genome v1.0 using Bismark (Krueger and Andrews, 2011) in the directional mode, employing the Bowtie2 mapper with pre-defined settings. Methylation data were extracted with Bismark_methylation_extractor, retaining only reads with a unique best alignment score. Methylation data were then analyzed in the Methylkit R package (Akalin et al., 2012).

### Estimation of Bisulfite Conversion Efficiency

We also mapped the filtered reads to chloroplast genomes (assumed to be completely unmethylated) in order to assess the bisulfite conversion efficiency. Reads from Recital, Langdon, Jx109, Jx87 and Lx109-C2 were aligned to the chloroplast genome of Chinese Spring (GeneBank ID NC_002762.1). Reads from 109×L-C2, 109×L-C4, Tauschii-109 and Tauschii-87 were mapped to the *Ae. tauschii* chloroplast genome (NC_022133.1). The conversion rate (Table S2) was then calculated following the formula: conversion rate=(number of converted cytosines/total number of analyzed cytosines)×100

### Data analysis

We further filtered out presumed PCR artifacts by removing bases with coverage above the 99.99th percentile, using the ‘filterByCoverage’ function in methylKit. Cytosine methylation was summarized using two different calculations and coverage thresholds, depending on the type of the analysis. For single-cytosine analyses (i.e. detection of differentially methylated cytosines), we only considered positions with a minimum of 10x coverage. For global methylation averages across all genes summarized per subgenome and sequence context, mean methylation level (Schultz et al., 2012) was calculated from all positions with ≥10x coverage. For region-based methylation analyses (i.e. DNA methylation scans along chromosomes, all TSS/SC-based analyses in relation to gene expression and differential methylation), weighted methylation level (Schultz et al., 2012) was calculated from all cytosines with coverage ≥3. We consider such summarization optimal for the region-based analyses, since it maximizes the number of evaluated cytosines in an unbiased way (a low-coverage site contributes proportionally less to the weighted methylation level compared to a high-coverage site). Fine-scale methylation around TSS/SCs was summarized in 10bp bins using the ScoreMatrixBin function from genomation package (Akalin et al., 2015).

Differentially methylated regions (DMRs) and differentially methylated cytosines (DMCs) were identified between hexaploid synthetics and parents (except for the synthetics Jx87-S5 and Jx109-S5, where BSseq data of the tetraploid parent are not available) using the ‘calculateDiffMeth’ function in methylKit that accounts for variation between replicates. It employs a logistic regression-based method with overdispersion correction and adjusts the p-values using the Benjamini-Hochberg method (FDR) (Akalin et al., 2012). Regions/cytosines with a methylation level difference greater than 25% and FDR <0.01 were considered differentially methylated. For the DMR identification, we employed a model-based method using the methseg function in methylKit, which divides the target regions into variable-length segments of homogeneous methylation levels (Akalin et al., 2012). Segments identified in the parents (Tauschii-109 and Langdon) were used to screen the parents and their derived synthetics.

## DATA AVAILABILITY STATEMENT

The capture-BSseq data produced and analyzed for the purpose of this study are available at the Sequence Read Archive (https://www.ncbi.nlm.nih.gov/sra) under the BioProject number PRJNA1247774.

## ACKNOWLEDGEMENTS

This work was partially supported by the INRAE BAP department (project Methylwheat) and by a PhD fellowship awarded to MB (jointly by I-SITE 20-25 and INRAE BAP department). We are thankful to Annaig Bouguennec for the production of wheat synthetics, and to Cecile Huneau for nucleic acid extractions. A part of the experiments (capture-BSseq library preparation) was performed at the PGTB (doi:10.15454/1.5572396583599417E12), with the help of Christophe Boury and Erwan Guichoux. We are grateful to the Mésocentre Clermont-Auvergne and AuBi (plateforme Auvergne Bioinformatique) platform of the Université Clermont Auvergne for their support, computing and storage resources.

## SHORT LEGENDS FOR SUPPORTING INFORMATION

Supporting_Information_1: Contains supplementary tables Table S1-S6.

Supporting_Information_2: Contains supplementary figures Figure S1-S11.

