## Supplementary Figures for "DNA methylation around transcription start sites not globally associated with transcription in the grain of natural and synthetic hexaploid wheat"

for

### Supplementary Figures S1-S11

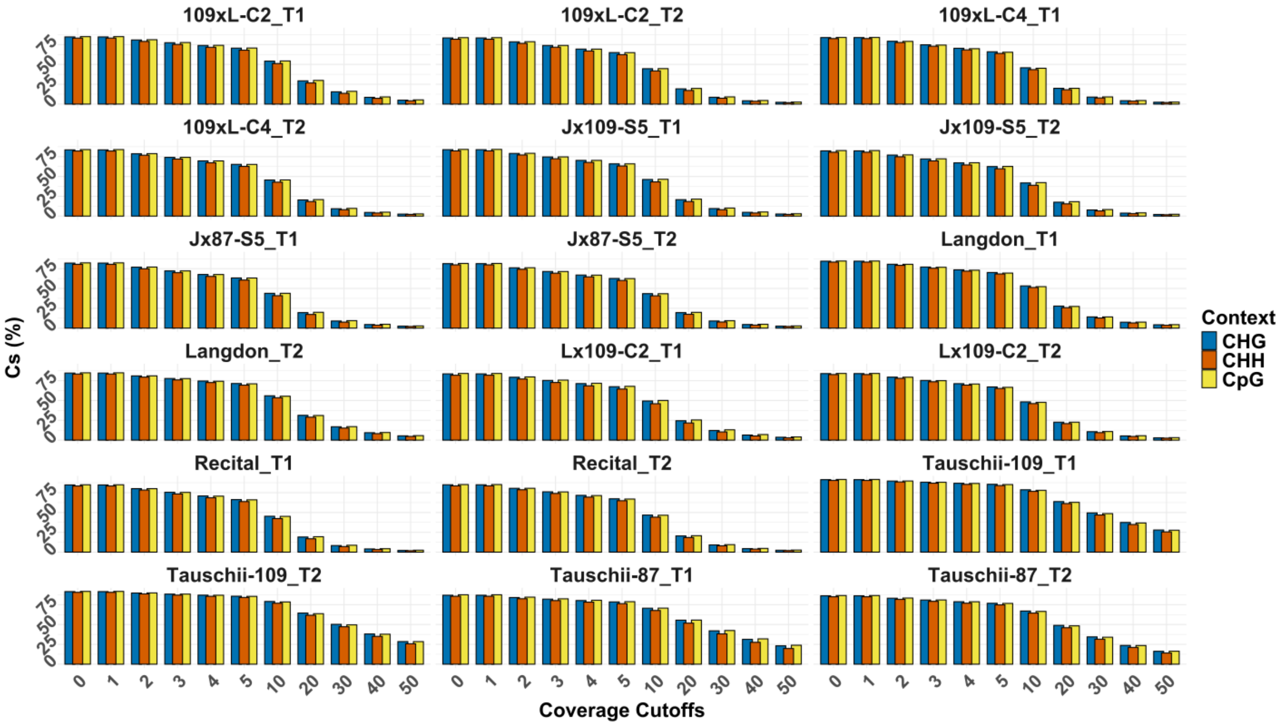


**Figure S1.** Coverage of target cytosine sites at various depth thresholds. Percentage of target cytosines covered by data at depth thresholds indicated on the x-axis. Each library was processed separately, with suffixes T1 and T2 indicating the biological replicates.

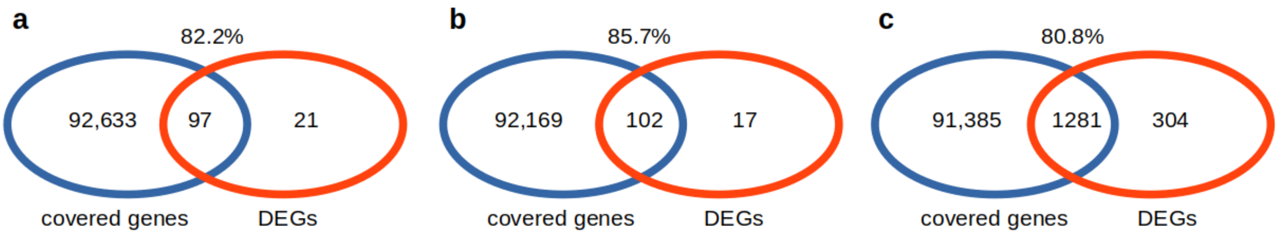


**Figure S2.** Coverage of DEGs identified in Banouh et al. (2023). Total number of high-confidence genes (Chinese Spring v1.1 annotation) covered at the 3x depth threshold, in relation to DEGs identified previously from the same samples and analyzed in this paper. (a) 109xL-C2; (b) 109xL-C4; (c) Lx109-C2.

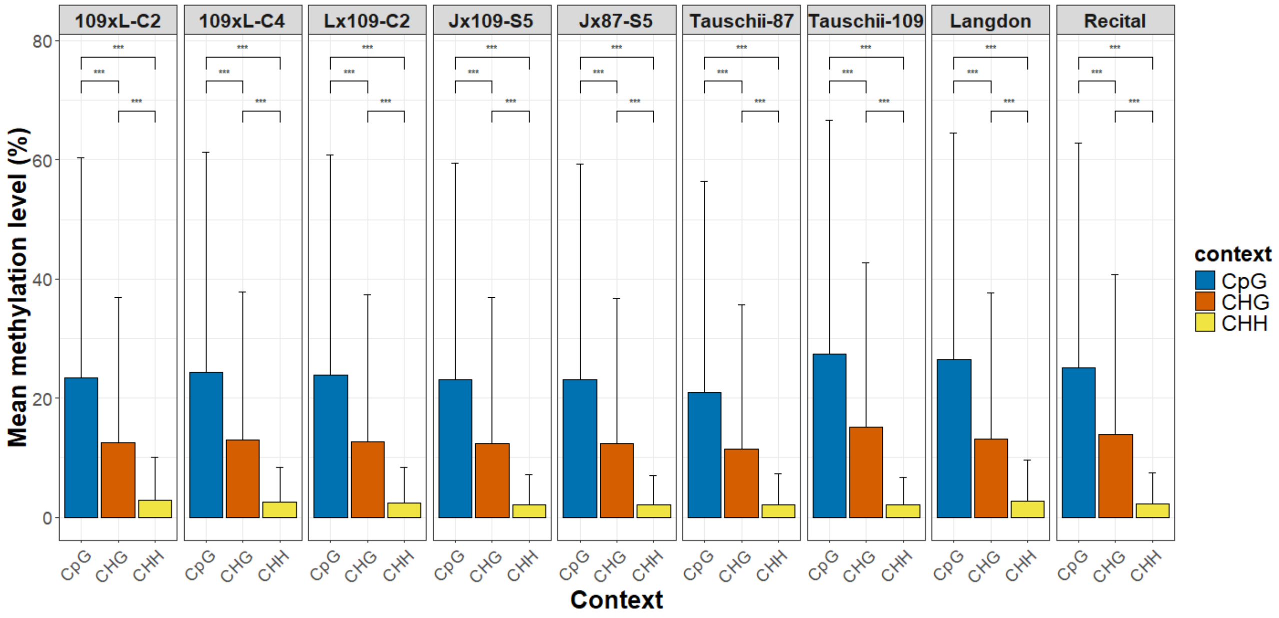


**Figure S3.** Mean methylation levels per genotype across different sequence contexts (CpG, CHG, CHH). Statistical significance (Wilcoxon test) is indicated with asterisks (*** $p < 0.001$).

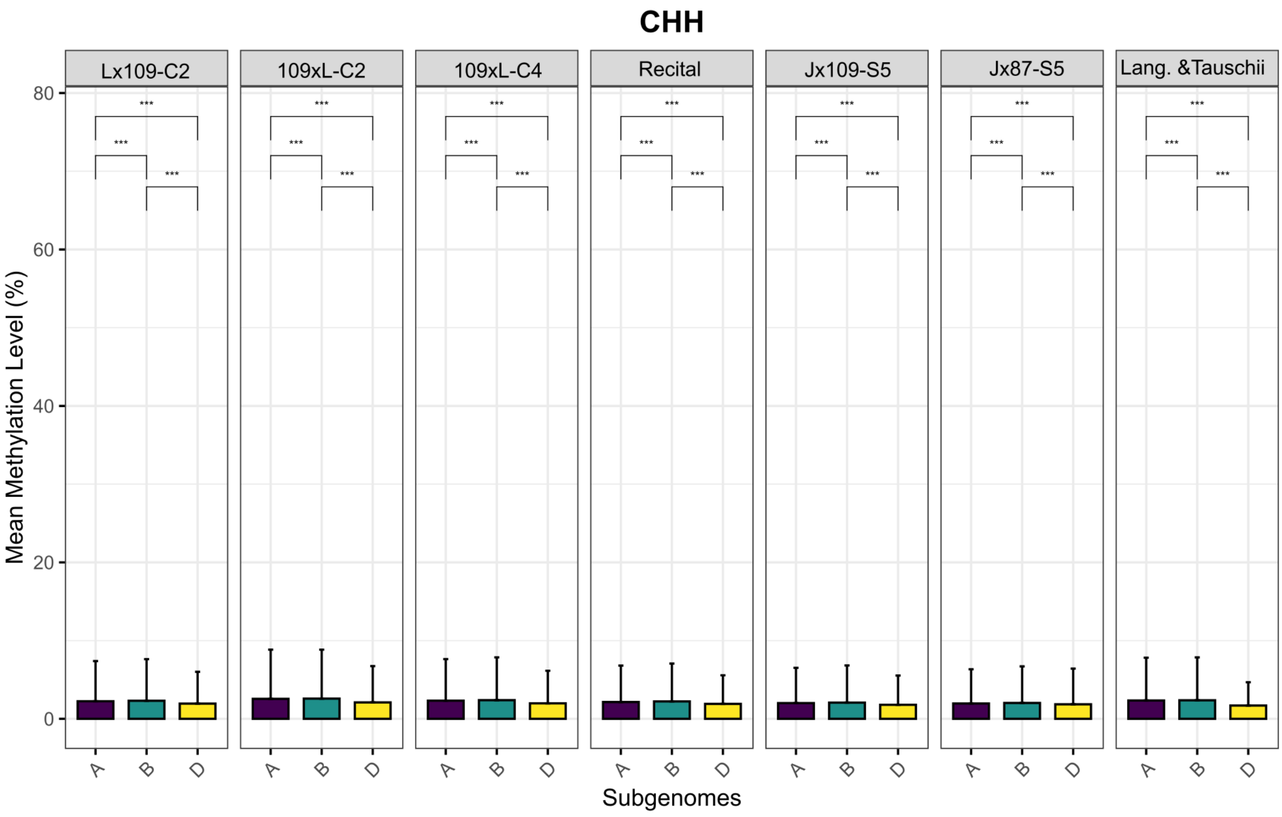


**Figure S4.** Mean Methylation Levels per subgenome (A, B and D) across genotypes in CHH contexts (*** $p < 0.001$). Error bars represent the standard deviation.

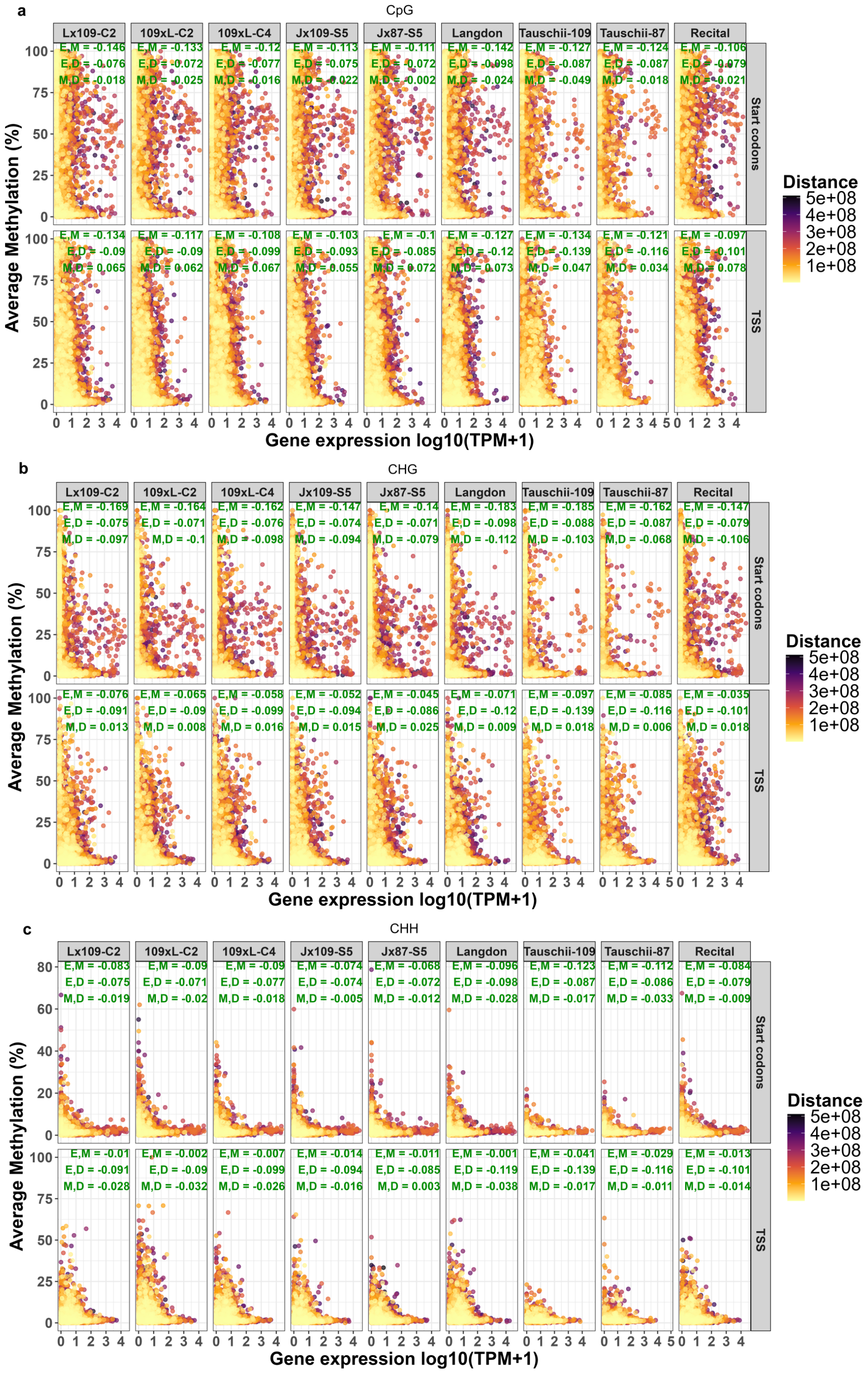


**Figure S5.** The relationship between transcription (x-axis), DNA methylation level (y-axis), and centromere distance (expressed with a color code). Pearson's correlation between gene expression and DNA methylation (E,M), gene expression and centromere distance (E,D), and DNA methylation and centromere distance (M,D) is show in green for each scatter plot. DNA methylation is shown separately for the CpG context (a), CHG context (b) and CHH context (c). For each context, the top panels show the SC-centered data set, while the bottom panels show TSS-centered data set.

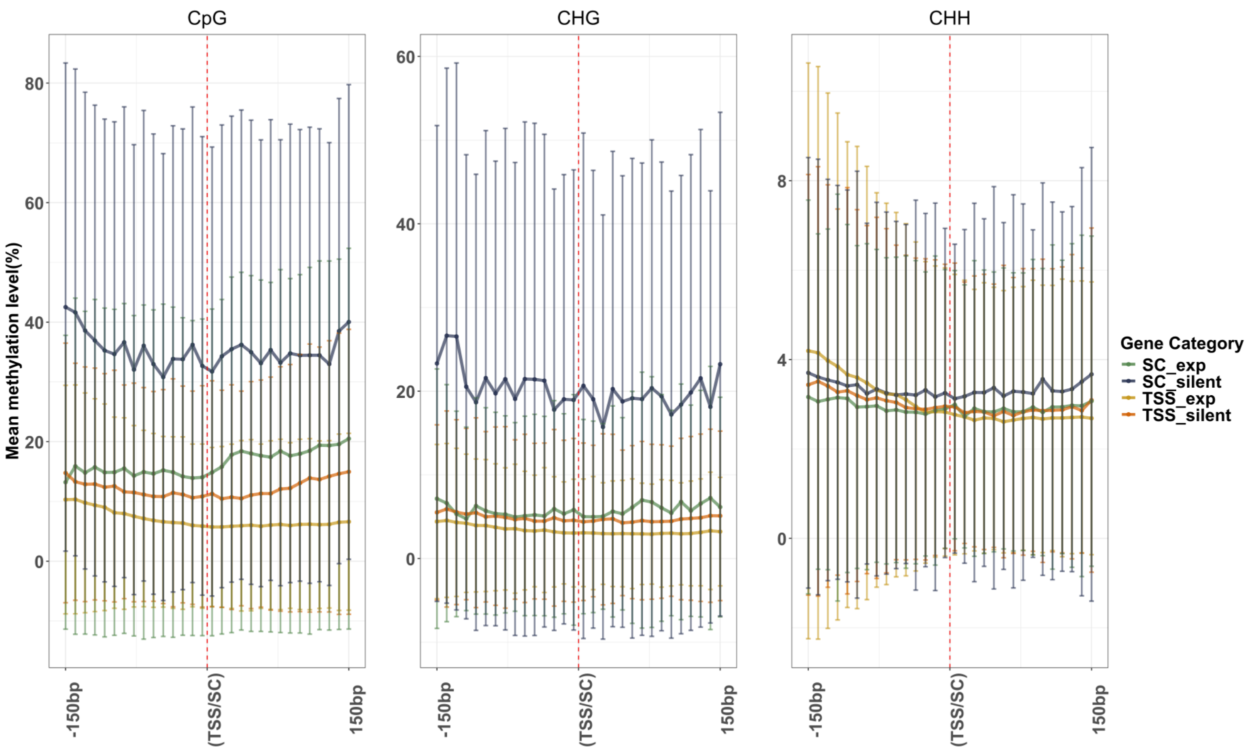


**Figure S6.** Mean methylation levels at ten-base pair resolutions around TSS and SC of all expressed ('exp') and silent genes of the natural allohexaploid Recital. The error bars represent standard deviations across genes, with large overlaps across the gene categories.

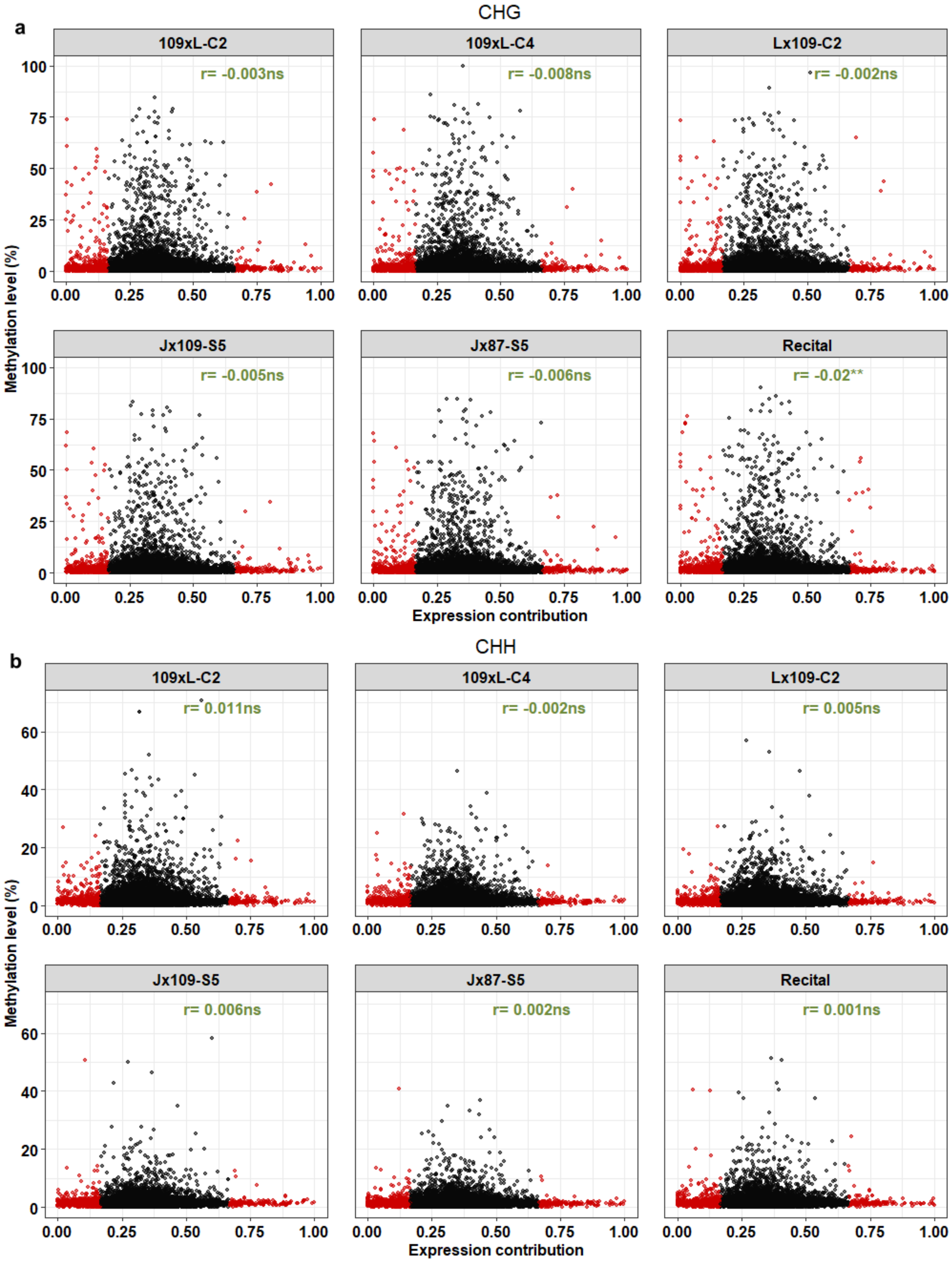


**Figure S7.** Scatter plots showing the relationship between methylation level and expression contribution to the triad. Suppressed and dominant genes are shown in red. (a) CHG context (b) CHH contexts. Pearson's correlation coefficient are shown for each sample in green (ns, $p$ >0.05; ** $p$ < 0.01).

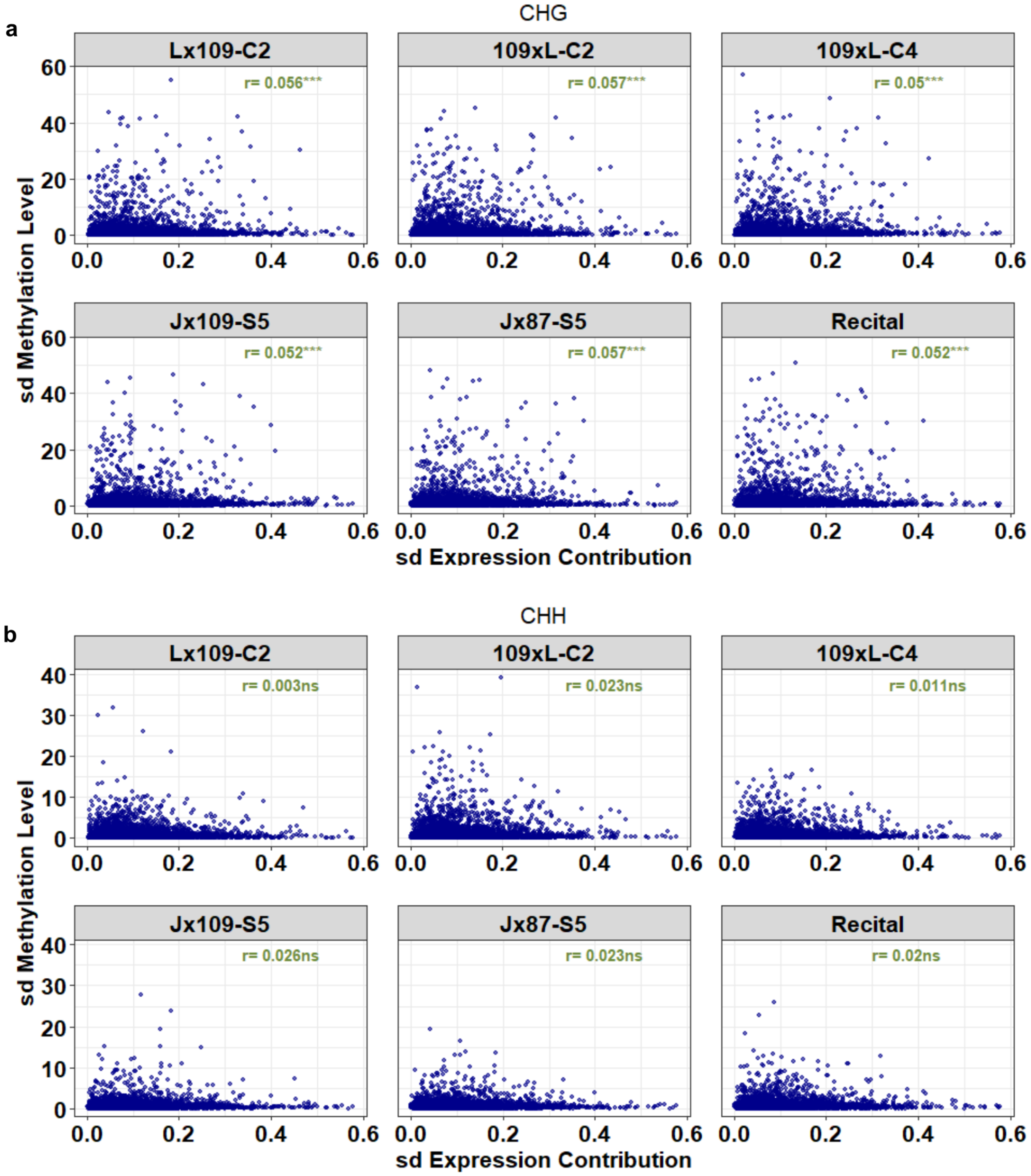


**Figure S8.** Scatter plots showing the correlation between the variance (sd) in expression contribution within triads, and the corresponding variance in methylation levels within triads in (a) CHG and (b) CHH contexts. Pearson's correlation coefficients are shown for each sample in green (ns, $p > 0.05$; *** $p < 0.001$).

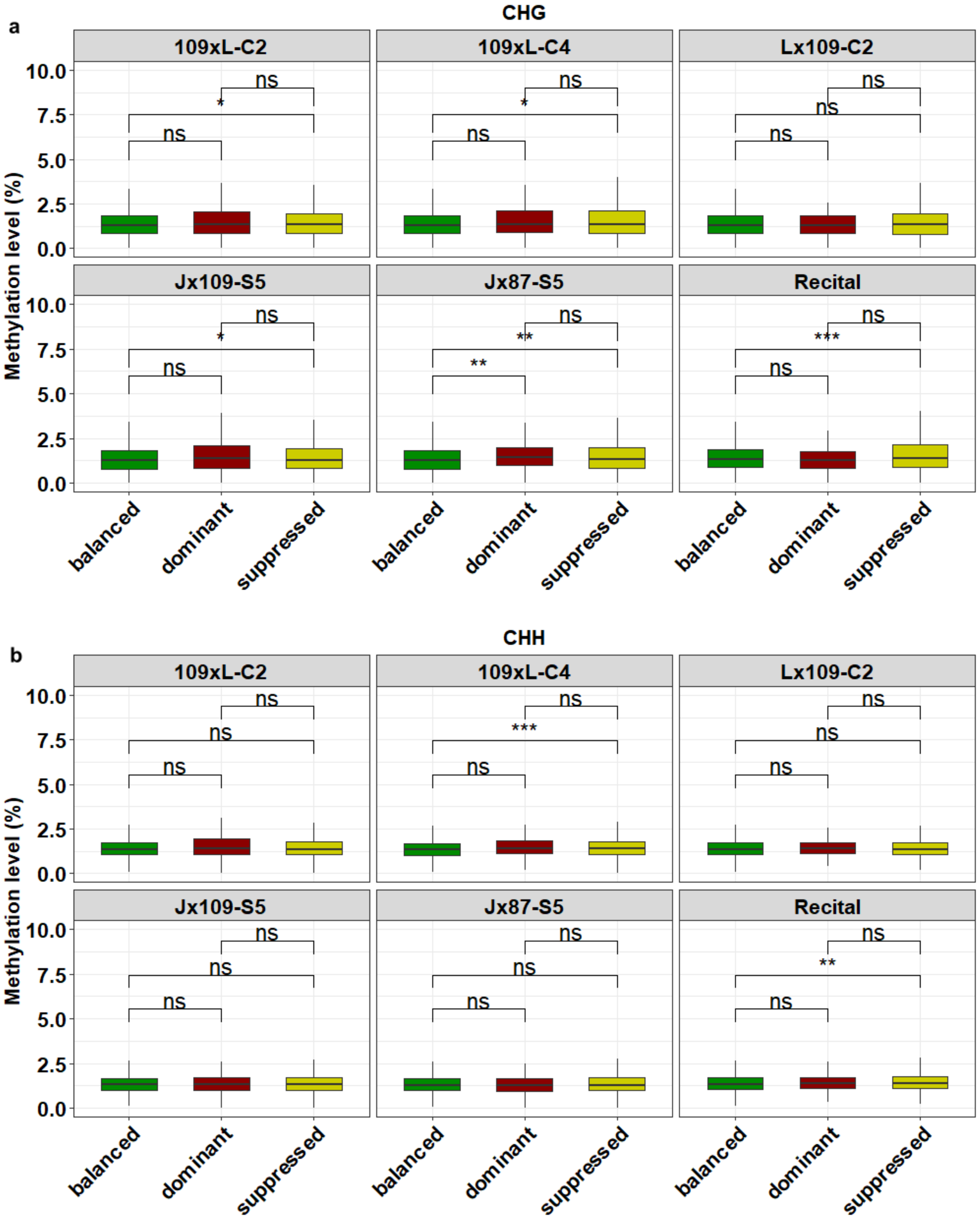


**Figure S9.** Comparisons of mean methylation between balanced, suppressed and dominant genes across samples with error bars representing the mean ± SD. Asterisks indicate significant differences based on the Wilcoxon test (ns = not significant, * $p < 0.05$, ** $p < 0.01$, *** $p < 0.001$).

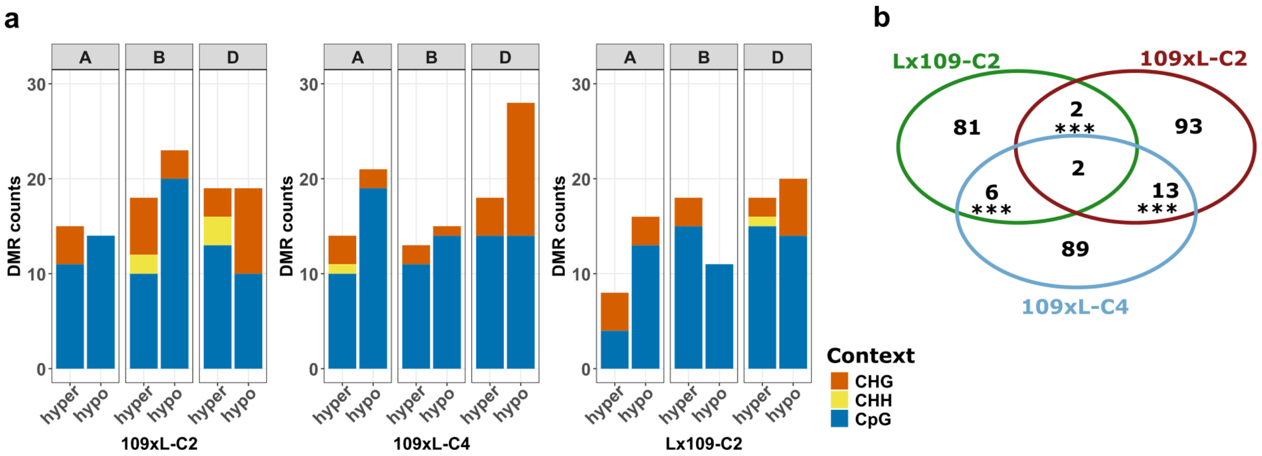


**Figure S10.** DMRs identification using 300bp windows as regions. (a) Counts of hyper/hypomethylated DMRs in different hexaploid synthetics across different subgenomes and different cytosine contexts. (b) Venn diagram showing the overlap of DMRs (considering all contexts) detected in different synthetic samples. Asterisks indicate the significance of the overlaps (*** $p < 0.001$).

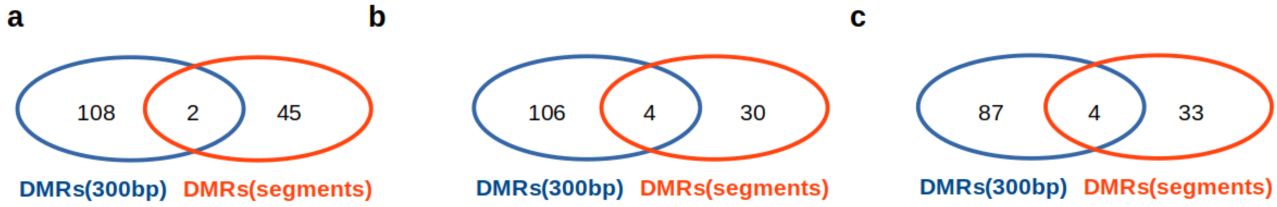


**Figure S11.** Venn diagrams showing the concordance (or lack thereof) of the two methods used for the DMR detection (considering all contexts). (a) 109xL-C2; (b) 109xL-C4; (c) Lx109-C2.
